# Targeted and Explorative Profiling of Kallikrein Proteases and Global Proteome Biology of Pancreatic Ductal Adenocarcinoma, Chronic Pancreatitis, and Normal Pancreas Highlights Disease-Specific Proteome Remodelling

**DOI:** 10.1101/2022.11.28.518142

**Authors:** Janina Werner, Patrick Bernhard, Miguel Cosenza-Contreras, Niko Pinter, Matthias Fahrner, Prama Pallavi, Johannes Eberhard, Peter Bronsert, Felix Rückert, Oliver Schilling

## Abstract

Pancreatic ductal adenocarcinoma (PDAC) represents one of the most aggressive and lethal malignancies worldwide with an urgent need for new diagnostic and therapeutic strategies. One major risk factor for PDAC is the pre-indication of chronic pancreatitis (CP), which represents highly inflammatory pancreatic tissue. Kallikreins (KLKs) are secreted serine proteases that play an important role in various cancers as components of the tumor microenvironment. Previous studies of KLKs in solid tumors largely relied on either transcriptomics or immunodetection. We present one of the first targeted mass spectrometry profiling of kallikrein proteases in PDAC, CP, and normal pancreas. We show that KLK6 and KLK10 are significantly upregulated in PDAC (n=14) but not in CP (n=7) when compared to normal pancreas (n=21), highlighting their specific intertwining with malignancy. Additional explorative proteome profiling identified 5936 proteins in our pancreatic cohort and observed disease-specific proteome rearrangements in PDAC and CP. As such, PDAC features an enriched proteome motif for extracellular matrix (ECM) and cell adhesion while there is depletion of mitochondrial energy metabolism proteins, reminiscent of the Warburg effect. Although often regarded as a PDAC hallmark, the ECM fingerprint was also observed in CP, alongside with a prototypical inflammatory proteome motif as well as with an increased wound healing process and proteolytic activity, thereby possibly illustrating tissue autolysis. Proteogenomic analysis based on publicly accessible data sources identified 112 PDAC-specific and 32 CP-specific single amino acid variants, which among others affect KRAS and ANKHD1. Our study emphasizes the diagnostic potential of kallikreins and provides novel insights into proteomic characteristics of PDAC and CP.

## Introduction

Pancreatic ductal adenocarcinoma (PDAC) represents the most common type of pancreatic cancer, accounting for more than 90 % of all pancreatic malignancies (Sarantis, Koustas, Papadimitropoulou, Papavassiliou, & Karamouzis, 2020). It is characterized by a highly aggressive and invasive growth, resulting in an abysmal 5-year survival rate of merely 3-15 % (Pereira et al., 2020). To date, surgical resection is the only option for cure (Casolino et al., 2021). However, the majority of patients receive palliative treatment, as most patients present with their disease at an already locally advanced or metastatic state (McGuigan et al., 2018). Although the protein carbohydrate antigen 19-9 (Ca 19-9), the only clinically accepted and routinely used biomarker for PDAC, serves as a vital follow-up parameter during cancer progression and response to treatment, it is insufficient for screening and diagnostic among the general population (Kim et al., 2004; Luo et al., 2021). This highlights the lack of diagnostic approaches for PDAC and illustrates the urgent need for the identification of novel biomarkers which enable early diagnosis and subsequent therapy.

Challenges in the early detection of pancreatic cancer further include identifying at-risk individuals among the general population who would particularly benefit from diagnostic biomarkers for PDAC (Singhi, Koay, Chari, & Maitra, 2019). One of these major risk factors for the development of PDAC represents chronic pancreatitis (CP), which is characterized by recurrent, highly inflammatory episodes in the pancreatic tissue and has been discussed to be a preliminary stage to PDAC (Alhobayb, Peravali, & Ashkar, 2021; Del Poggetto et al., 2021; Vujasinovic et al., 2020).

Human kallikreins (KLKs) comprise a family of 15 secreted serine proteases. For PDAC, KLK 6 and 10 have been suggested to represent new diagnostic and/or prognostic biomarkers for pancreatic cancer (X. Y. Cao et al., 2018; Hustinx et al., 2004; Iacobuzio-Donahue et al., 2003; Ruckert et al., 2008; Yousef, Borgono, White, et al., 2004). Moreover, KLK3, known as prostate-specific antigen (PSA), is already of great clinical value as a routinely applied diagnostic marker and follow-up parameter in prostate cancer (Hong, 2014). Some kallikreins have been proposed as therapeutic targets for pancreatic cancer, as elevated levels of KLK6, KLK7, KLK8, KLK10, and KLK11 in PDAC were correlated with a significantly worse overall patient survival and are further suggested to play a crucial role within pancreatic cancer development (Candido et al., 2021; X. Y. Cao et al., 2018; Iakovlev, Siegel, Tsao, & Haun, 2012; Ruckert et al., 2008). Various pro-tumorigenic mechanistic routes have been proposed such as facilitating cancer cell migration by cleavage of cell adhesion and extracellular matrix proteins, suggesting KLKs as important factors in modulating the tumor microenvironment (S. K. Johnson, Ramani, Hennings, & Haun, 2007; Klucky et al., 2007; Magklara et al., 2003; Srinivasan, Kryza, Batra, & Clements, 2022). KLK6 levels were found to be significantly elevated in invasive tumor areas compared with non-invasive tumor areas in PDAC tissue sections (Candido et al., 2021). Besides the upregulation of kallikreins in PDAC tissues, multiple kallikreins, including KLK5, KLK6, KLK7 and KLK10, were also found to be elevated in cell-conditioned medium of the pancreatic cancer cell line MiaPaCa-2 (Shaw & Diamandis, 2008). However, currently available data on KLK secretion or expression in pancreatic cancer cell lines almost exclusively originates from transcriptomic data or immunodetection such as Western blot analysis, immunohistochemistry, or ELISA, which are often limited to few targets per study (X. Y. Cao et al., 2018; Ruckert et al., 2008; Shaw & Diamandis, 2008).

Gene or protein expression profiling approaches are increasingly used to gain deep insight into pathological traits pancreatic malignancies, as for example moffit and CPTAC. In this study, we applied targeted mass spectrometry (MS), namely parallel reaction monitoring (PRM) to investigate KLK expression in PDAC, ampullary cancer (AMPAC), CP, and normal non-malignant adjacent pancreas. To our knowledge, this is one of the first reports using targeted mass spectrometry for the detection of KLK proteins in these disease conditions. Furthermore, using exploratory proteomics, we performed proteome profiling of PDAC, AMPAC, CP, and normal non-malignant adjacent pancreas to gain better insight into the disease-related proteome biology.

## Materials and Methods

### Cell Culture and Preparation of Cell-conditioned medium (CCM)

The human pancreatic cancer cell line MiaPaCa-2 was cultured in Dulbecco’s modified Eagle’s medium (DMEM, high glucose, GlutaMAX™ supplement; Gibco, Thermo Fisher Scientific), while the human pancreatic cancer cell line AsPC-1 (passage 11) was cultured in RPMI-1640 medium (L-glutamine supplement; Gibco, Thermo Fisher Scientific), both supplemented with 10 % (v/v) Fetal bovine serum (FBS, Gibco, Thermo Fisher Scientific), 100 U/mL penicillin and 100 μg/mL streptomycin at 37 °C under a humidified atmosphere containing 5 % CO_2_. The human primary pancreatic cancer cell lines PaCaDD-159, PaCaDD-165 and MaPaC-107 were cultured in a 2:1 mixture of supplemented DMEM as described above and Keratinocyte-Serum-free medium (KSFM, Thermo Fisher Scientific) supplemented with 2.5 μg Epidermal Growth Factor (EGF, Human Recombinant) and 25 mg Bovine Pituitary Extract (BPE) at 37 °C under a humidified atmosphere containing 5 % CO_2_. All cell lines were maintained and subcultured at a cell confluency between 50-90 % and were regularly tested for mycoplasma contaminations (Eurofins Genomics).

For the preparation of cell-conditioned medium (CCM), cells were cultured in 145 mm cell culture dishes (Greine Bio-One) and expanded until 70-90 % confluency. Subsequently, cells were incubated for 24 h in serum-free medium (SFM, Gibco, Thermo Fisher Scientific) supplemented with 100 U/mL penicillin, 100 μg/mL streptomycin, 1 % (v/v) Minimum Essential Medium (MEM) solution and 1 % (v/v) Non-essential Amino Acid solution. Resulting cell-conditioned medium (CCM) was separated and supplemented with 10 mM EDTA and 0.1 mM PMSF followed by centrifugation (4000 rpm, 10 min) and storage of the clear supernatant at - 80 °C until further processing. The corresponding cell number of each CCM dish was determined by harvesting cells using TrypLE™ Express Enzyme trypsin-EDTA solution (Gibco, Thermo Fisher Scientific) and counting cells by using 0.4 % (v/v) Trypan blue staining solution (NanoEntek) and an automated cell counter (Eve Automatic Cell Counter, NanoEnTek).

### Cohort Information

The patient cohort included formalin-fixed paraffin-embedded (FFPE) tissues of PDAC (n=14), chronic pancreatitis (n=7), non-malignant adjacent pancreas (n=11), normal pancreas (n=5) and ampullary cancer (n=3), which were kindly provided by the surgical department of the University of Heidelberg, Medical Faculty Mannheim (PDAC, chronic pancreatitis and non-malignant adjacent pancreas, ampullary cancer), and the University of Freiburg, Institute for Surgical Pathology (normal pancreas). The latter samples were obtained from the pancreas of multiple organ donors (median age 63 years; range 43 ± 84). FFPE blocks of each tissue unit were cut at different thicknesses ranging from 1.5 - 10 μm and dried overnight at 60 °C by using a microtome (HM355 S, Thermo Fisher Scientific). All tissue slices were routinely stained with hematoxylin and eosin (H&E) using the Dako Coverstainer (Agilent Technologies) and were reviewed by an experienced pathologist. The area of interest was then marked on the tissue slices; regions that were not marked on the slide were excluded from further experiments. Written informed consent was obtained from all patients before inclusion into our study. Ethics approval was obtained by local authorities of the Ethics Committee of the University Medical Center Freiburg (file number: 367/14) and the Ethics Committee of the University Medical Center Heidelberg, Faculty of Mannheim (file number: 2012-293N-MA).

### Protein Extraction and Proteomic Sample Preparation

For sample preparation of cell-conditioned medium, 5 mL CCM of each cell line was transferred to a 15 mL falcon and was supplemented with 1 μg Avidin (Sigma) as internal standard and with 0.2 % (v/v) SDS, 35 mM TRIS pH 8.5 and 9 U/mL benzonase for protein extraction, respectively. Samples were incubated for 15 min at ambient temperature before performing cysteine reduction using 2 mM tris(2-carboxyethyl)phosphin-hydrochlorid (TCEP, 30 min, 80 °C water bath) and subsequent alkylation using 5 mM iodoacetamide (30 min, room temperature, in the dark). Sample pH was adjusted to pH 7 by adding 83 mM 2-Morpholinoethanesulfonic acid monohydrate (MES) before performing protein enrichment and SDS removal by using a previously published sp3-bead protocol (Hughes et al., 2019). In short, sp3-beads (Sera-Mag SpeedBeads, GE Healthcare) were added to a final concentration of 0.2 μg/μL before adding acetonitrile to a final proportion of 60 % (v/v) and incubating for 10 min at room temperature to allow protein binding. Beads were pelleted by centrifugation (800 g, 8 min), resuspended in 80 % (v/v) ethanol and transferred to Eppendorf tubes to allow bead separation using a magnetic rack. Beads were successively washed with 80 % (v/v) ethanol and acetonitrile before air drying beads. Protein-coupled beads were resuspended in 0.1 M HEPES pH 8.0 containing 0.1 % (v/v) of an acid-labile surfactant (sodium 3-[(2-methyl-2-undecyl-1,3-dioxolan-4-yl)methoxy]-1-propanesulfonate) before eluting proteins by sonication (10 min) and heating (95 °C, 10 min). Beads were removed using the magnetic rack only using the clear supernatant for further processing. Protein concentration of each eluate was determined by using the BCA assay (Thermo Fisher Scientific) before subjecting 10-70 μg protein amount to in-solution tryptic digestion. Proteolytic digestion was performed by adding Lysyl Endopeptidase (LysC, Wako Chemicals) in a protease:protein ratio of 1:50 (w/w) and incubating for 2 hours at 42 °C, before adding trypsin (Promega) in a protease:protein ratio of 1:50 (w/w) and further incubate samples for 19 hours at 37 °C. Digestion was stopped by acidification (2 % (v/v) TFA), before incubating (37 °C, 30 min) and centrifuging (20,000 g, 5 min) the samples in order to precipitate acid-labile surfactant. Clear supernatant was used for peptide desalting using iST C18 mixed phase cartridges (PreOmics, Martinsried, Germany) according to manufacturer’s instructions. After determining the peptide concentration via the BCA assay (Thermo Fisher Scientific), eluates were vacuum dried and stored at – 80 °C until MS measurement.

For sample preparation of formalin-fixed, paraffin-embedded (FFPE) samples, tissue slices were deparaffinized by successively incubating in increasing concentrations of xylene followed by decreasing concentrations of ethanol (xylol 4x, alcohol absolute, alcohol 96 %, alcohol 70 %, aqua distilled 2x). Samples (volume ranging from 0.3-0.6 mm^3^) were macrodissected into Eppendorf tubes and stored at - 80 °C until further processing. For protein extraction, tissue samples were incubated with detergent-containing buffer (0.1 M HEPES pH 8.0, 0.1 % (v/v) SDS) before performing heat-induced antigen retrieval (2 h at 95 °C) followed by ultrasonication (Bioruptor, 10 cycles, 45/15 sec on/off time, high intensity) and centrifugation (21.000g, 10 min), thereupon only using the clear supernatant. The protein concentration was determined by using the BCA assay (Thermo Fisher Scientific) before subjecting 100 μg protein amount to in-solution proteomic sample preparation including cysteine reduction with 5 mM DTT (37 °C, 30min), subsequent alkylation with 15 mM iodoacetamide (30 min, room temperature, in the dark) and protein enrichment and SDS removal by using sp3-beads like described above. On-bead proteolytic digest was performed by adding Lysyl Endopeptidase (LysC, Wako Chemicals) in a protease:protein ratio of 1:100 (w/w) and incubating for 2 hours at 42 °C, before adding trypsin (Promega) in a protease:protein ratio of 1:50 (w/w) and further incubate samples for 19 hours at 37 °C. Post-digestion processing was performed like described above.

### LC-MS/MS measurement

For SRM measurement, vacuum dried peptides were resolubilized in 0.1 % (v/v) formic acid to a final concentration of 0.2 μg/μL, sonicated for 5 min and centrifuged at 20,000 g for 10 min before transferring the supernatant to the measurement tube. 1 μg of each sample, together with 200 fmol of indexed retention time (iRT) peptides and 100 fmol of selected isotope-labelled heavy reference peptides, were analyzed using a nanoflow liquid chromatography (LC) system, Proxeon Easy-nLC II (Thermo Fisher Scientific) equipped with a trapping column (Fused Silica Capillary, 5 cm length, 100 μm inner diameter (ID), VICI Jour, Schenkon, Switzerland) and an analytical column (Selfpack PicoFrit column, 35 cm length, 75 μm ID, New Objective, Woburn, MA) both in-house packed with C18 particles (Dr. Maisch, ReproSil-Pur 120 C18-AQ, 3 μm C18 particle size, 120 Å pore size). Samples were trapped at 220 bars with 100 % Buffer A (0.1 % (v/v) formic acid) and separated using a flow rate of 250 nL/min with the analytical column temperature set to 60 °C. A 83-min multistep gradient of 8 % to 56 % (v/v) buffer B (50 % (v/v) ACN in 0.1 % (v/v) formic acid in H_2_O) in buffer A was used for separation followed by washing (100 % B) and reconditioning of the column to 5 % B (Table S1 for detailed gradient overview). The Easy-nLC II system was coupled online to a TSQ Vantage (Thermo Fisher Scientific) triple quadrupole mass spectrometer via a Nanopray Flex Ionsource (Thermo Fisher Scientific) with an applied voltage of 2.5 kV for electrospray ionization. The analytical PicoFrit column contained an integrated uncoated pre-cut emitter (Silica Tip, 10 μm tip inner diameter, New Objective). The mass spectrometer was operated in selected reaction monitoring (SRM) mode with scheduled acquisition. SRM acquisition was performed with Q1 and Q3 operated at unit resolution (0.70 m/z half maximum peak width) with 449 transitions and a fixed dwell time of 15 ms per transition. Collision energies (CEs) were calculated in Skyline according to the formulas: CE = 0.03 x m/z + 2.905 and CE = 0.038 x m/z + 2.281 (m/z, mass-to-charge ratio of the precursor ion) for doubly and triply charged precursor ions, respectively. Samples were measured in randomized sample order.

For PRM measurement, vacuum dried peptides were resolubilized like described above. 800 ng of each sample together with 100 fmol of iRT peptides and 100 fmol of selected isotope-labeled heavy reference peptides, were analyzed using a nanoflow LC system, Easy-nLC 1000 (Thermo Fisher Scientific) equipped with a trapping column (50 cm μPac^TM^ trapping column, Thermo Fisher Scientific) and an analytical column (200 cm μPac^TM^ analytical column, Thermo Fisher Scientific) tempered to 45 °C. Samples were trapped at 250 bars with 100% buffer A (0.1 % v/v formic acid) and separated using a dynamic flow rate of 350-700 nL/min. A 120-min multistep gradient of 8 % to 55 % buffer B (80 % v/v acetonitrile, 0.1 % v/v formic acid) in buffer A was used for separation, followed by washing (100 % B) and reconditioning of the column to 8 % B (Table S2 for detailed gradient overview). The Easy-nLC 1000 system was coupled online to a Q-Exactive plus (Thermo Fisher Scientific) mass spectrometer via a Nanospray Flex Ionsource (Thermo Fisher Scientific) with an applied voltage of 2.1 kV for electrospray ionization. The analytical column was coupled to a pulled, uncoated ESI emitter (10 μm tip inner diameter, 20 μm inner diameter, 7 cm length, CoAnn Technologies) via a μPacTM Flex iON Connect ESI-MS interface (PharmaFluidics). The mass spectrometer was operated in unscheduled parallel reaction monitoring (PRM) acquisition mode. Therefore, MS2 scans (1 μscan) of doubly-charged precursor ions were performed with an isolation window size of 1.2 m/z, MS2 resolution was set to 35’000, automatic gain control (AGC) to 3e6 and maximum injection time was set to 150 ms using stepped normalized collision energy (NCE) of 25 and 30 for fragmentation. Samples were measured in randomized sample order.

For DIA measurement, vacuum dried peptides were resolubilized like described above. 800 ng of each sample together with 100 fmol of iRT peptides, were analyzed using the same nanoflow LC and mass spectrometer system as described for PRM measurements.

For the individual DIA-samples, the mass spectrometer was operated in data independent acquisition (DIA) mode. Therefore, two cycles of 24 m/z broad windows ranging from 400 to 1000 m/z with a 50 % shift between the cycles (staggered window scheme) (B. C. Searle et al., 2018) was used, leading to an overall isolation window size of 12 m/z. MS2 resolution was set to 17,500, automatic gain control (AGC) to 1e6 and maximum injection time was set to 80 ms using stepped NCE of 25 and 30. After 25 consecutive MS2 scans, a MS1 survey scan was triggered covering the same range, but with a resolution of 70,000, an AGC target of 3e6 and maximum injection time was set to 50 ms. Samples were measured in randomized sample order.

For GPF-library measurements to generate an experiment-specific database, similar sample amounts were mixed to generate a masterpool sample, which was subsequently measured. The same settings were applied with the exception that the total mass scan range of 400 to 1,000 m/z has been subdivided into 6 individual measurements, each covering a mass scan range of 100 m/z with an overall isolation window size of 4 m/z.

### Proteomic Data Analysis

For SRM and PRM data, proteomic data analysis was performed using the open source software tool Skyline (version 19.1) (MacLean et al., 2010). All integrated peaks were manually inspected to ensure correct peak detection and integration. The resulting peak areas of KLK proteins were successively normalized to both the internal standard avidin and the total cell number the CCM derived from. For statistical evaluation, a two-sample t-test was performed using Benjamini-Hochberg for multiple testing correction. Resulting adjusted p-values and corresponding fold changes were visualized as volcano plots, with proteins revealing a significant abundance change (adjusted p-value ≤ 0.05) together with an absolute log2 fold change ≥ 1 (corresponds to 2-fold abundance change) were considered as differentially expressed.

For DIA data analysis, DIA-NN (1.7.12) was used with recommended settings (Demichev, Messner, Vernardis, Lilley, & Ralser, 2020). Peptide identification was performed using a human proteome database containing reviewed UniProt sequences without isoforms downloaded from Uniprot on 14th June 2021 (20856 entries). This database was refined using the GPF-measurements. For library refinement, deep learning-based spectra and RTs prediction was activated. Tryptic cleavage specificity (Trypsin/P) with 1 missed cleavage was applied, while setting an allowed peptide length between 7-30 amino acids. The N-terminal M excision, C carbamidomethylation, retention time (RT) profiling and RT-dependent cross-normalization options were enabled. Precursor FDR was set to 1.0 %, while decoys of the database entries were automatically created by DIA-NN. For the in silico predicted library search, the same settings were applied with exception of deactivated deep learning-based spectra and RTs prediction option.

The DIA-NN output was further processed in R (V 4.1.0) with RStudio (V 1.3.1093) as an integrated development environment. After filtering for unique proteins, log2-transformation and median centering was performed using the DIANN package (V 1.0.1) (Demichev et al., 2020). For sparsity reduction, the dataset was filtered for proteins with at least 80 % valid values per protein, per condition before performing ComBat batch correction by using the sva (V 3.40.0) package (W. E. Johnson, Li, & Rabinovic, 2007).

Partial least squares discriminant analysis (PLS-DA) and extraction of protein_representatives was performed by using the mixOmics package (V 6.16.3). Cluster of proteins that are consistently co-expressed were identified by using the Clust algorithm (V 1.10.10) (Clust Citation) in Python (V 3.8.9) using Z-score normalization prior to k-means clustering with seed number k=8 and tightness weight = 0.5. Differential expression analysis was performed using the limma (V 3.48.3) package. Gene Ontology annotation and gene enrichment analysis of protein subsets were performed using the topgo (V 2.44.0), clusterprofiler (V 4.0.5) and ReactomePA (V 1.36.0) packages.

For Semi-Tryptic Peptide Analysis of DIA data, peptide identification was performed like described above but using a human proteome database already containing semi-trpyptic peptides (10514442 entries) deriving from the abovementioned human database downloaded from Uniprot on 14th June 2021. After additional peptide-to-protein mapping by using an_in-house developed publicly available R-script (publicly available via repository: pep2prot (Version 1.2), https://github.com/npinter/pep2prot) and subsequent log2-transformation and median centering like described above, identified peptides were annotated as tryptic or semi-tryptic peptides by using another in-house developed publicly available R-script (publicly available via repository: Fragterminomics (Version 1.0.0), https://github.com/MiguelCos/Fragterminomics).

For proteogenomic analysis, the open-source and free-to-use Galaxy platform (Afgan et al., 2018) was used on the European Galaxy server (https://usegalaxy.eu). The tools for generating a proteogenomic FASTA database are connected within a Galaxy workflow that enables a standardized and parallel high-throughput analysis combined with file collections. Selected Paired-end RNA-seq data from the Sequence Read Archive (SRA) database (https://www.ncbi.nlm.nih.gov/sra) were used as input for the Galaxy workflow. To validate the quality of the provided input files, quality control was performed using FastQC (v0.73) and MultiQC (v1.11). First, raw RNA-seq reads were mapped to the human reference genome GRCh38 using HISAT2 (v2.2.1). Subsequently, the output of HISAT2 was deployed for variant calling using FreeBayes (v1.3.1). The resulting variant call file (VCF) and the mapped RNA-seq reads (BAM) were submitted to CustomProDB (v1.22.0) for generation of the proteogenomic FASTA database containing splice isoforms, single amino acid variants (SAAVs) as well as insertion and deletion (InDels) mutations. Further post-processing using Galaxy’s standard text manipulation tools yield the final proteoform database containing single amino acid variants (SAAVs). Subsequently, this database was processed locally with Python scripts to generate tryptic variant peptides (10806 entries). The abovementioned canonical human proteome database (downloaded from Uniprot on 14th June 2021; 20856 entries) was subsetted (subFASTA proteome) with the protein identifiers (UniProt IDs) from the initial DIA search and utilized for further filtering of those tryptic SAAV peptides to obtain a unique and peptide-centric variant database. The subFASTA proteome (6159 entries) and the generated SAAV database were merged and applied as a reference proteome for the variant peptide search in DIA-NN. Subsequent peptide identification from the acquired DIA data was performed like described above but without refinement using DIA-NN and the combined database (16965 entries).

## Results and Discussion

### KLK Detection in Cell-Conditioned Medium of Pancreatic Cancer Cell Lines

As a proof-of-concept system for mass spectrometric detection of secreted kallikrein proteases, we chose cell-conditioned medium (CCM) of cultured pancreatic cancer cells. Different human pancreatic cancer cell lines (MiaPaCa-2, AsPC-1, PaCaDD-165, PaCaDD-159 and MaPaC-107; Table 1) were maintained by standard cell culture before incubating in serum-free medium (SFM) to prepare CCM. Mass spectrometric analysis of this CCM by using selected reaction monitoring (SRM) enabled the targeted detection of secreted kallikreins from the respective cell lines. We probed all 15 human KLKs with a panel comprising 41 peptides, leading to the robust detection of KLK3, KLK5, KLK6, KLK7 and KLK10 in the considered cell lines (Fig. 1). Further KLKs remained below detection.

**Fig. 1:**
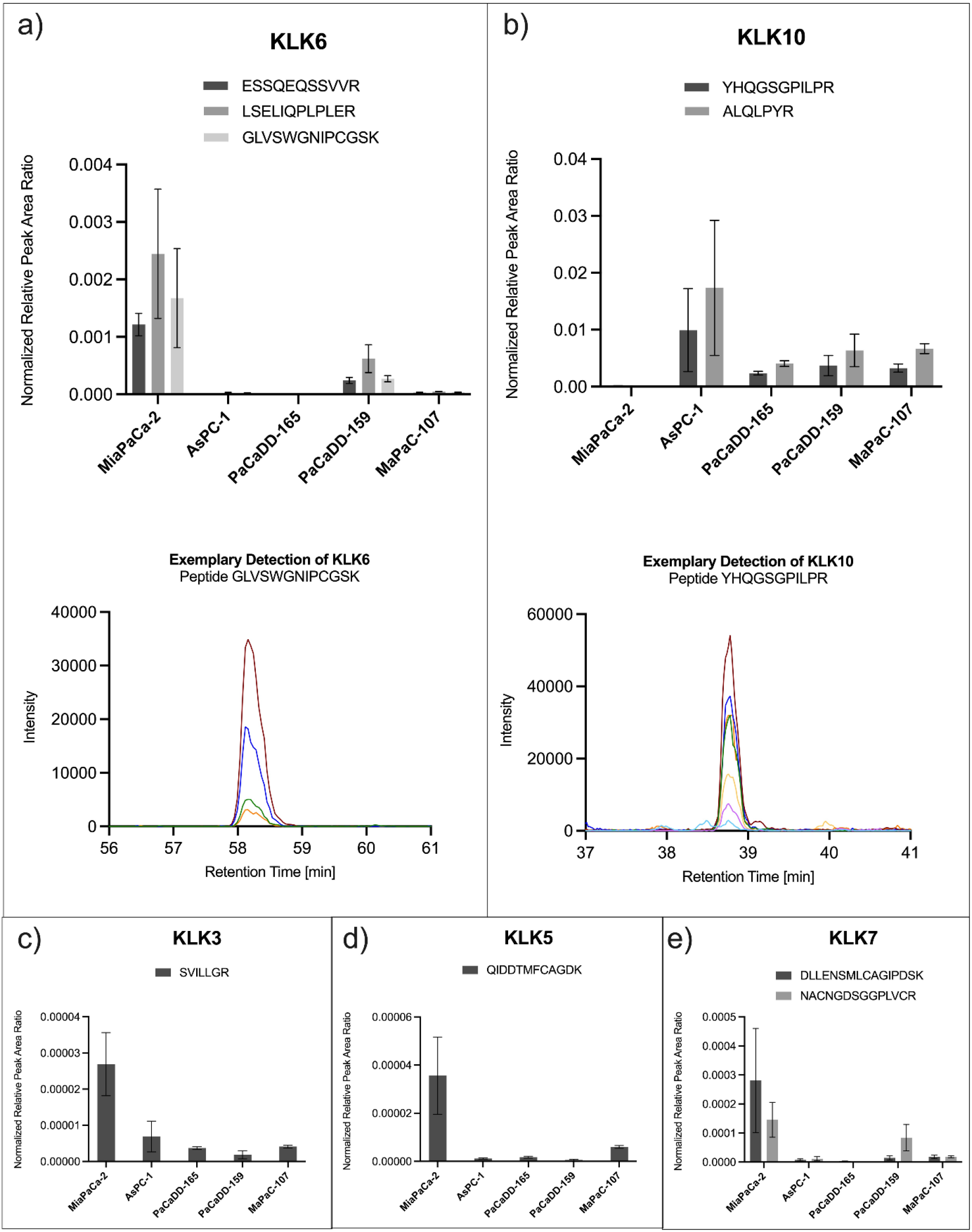
Targeted detection of secreted kallikrein proteins in cell-conditioned medium of pancreatic cancer cell lines by selected reaction monitoring. Serum-free, cell-conditioned medium of five pancreatic cancer cell lines (MiaPaCa-2, AsPC-1, PaCaDD-165, PaCaDD-159, MaPaC-107) were analyzed in a randomized sample order using selected reaction monitoring (SRM) to enable targeted detection and comparison of secreted kallikreins. The assay allowed for the detection of (a) KLK6 with three peptides and (b) KLK10 with two peptides, each of them with several fragments, exemplarily depicted below as co-eluting chromatographic peaks in different colors. Furthermore, (c) KLK3, (d) KLK5 and (e) KLK7 was detected with one or two peptides. Detected peak areas were successively normalized to avidin as internal standard and to the total cell number the CCM derived from. Bar plots show normalized peak area ratios after 2-step normalization for each peptide in each of the investigated cell lines. Error bars correspond to standard deviation.

**Table 1:**
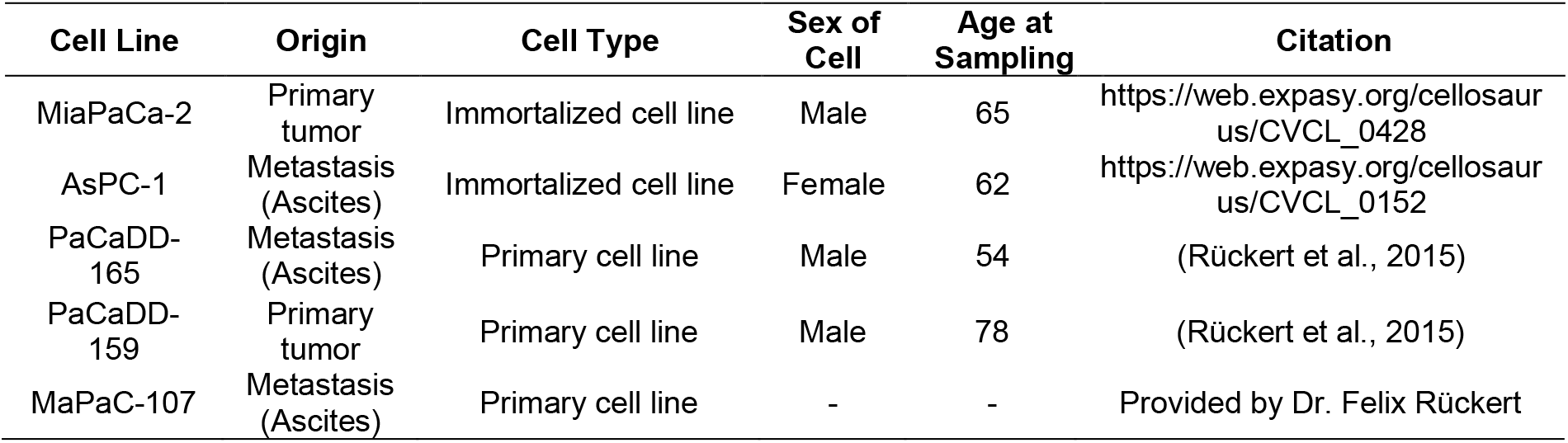
Characteristics of included pancreatic cancer cell lines. The used cell lines vary in their origin, cell type, sex, and age at sampling.

| Cell Line | Origin | Cell Type | Sex of Cell | Age at Sampling | Citation |
| --- | --- | --- | --- | --- | --- |
| MiaPaCa-2 | Primary tumor | Immortalized cell line | Male | 65 | https://web.expasy.org/cellosaurus/CVCL_0428 |
| AsPC-1 | Metastasis (Ascites) | Immortalized cell line | Female | 62 | https://web.expasy.org/cellosaurus/CVCL_0152 |
| PaCaDD-165 | Metastasis (Ascites) | Primary cell line | Male | 54 | (Rückert et al., 2015) |
| PaCaDD-159 | Primary tumor | Primary cell line | Male | 78 | (Rückert et al., 2015) |
| MaPaC-107 | Metastasis (Ascites) | Primary cell line | - | - | Provided by Dr. Felix Rückert |

We have refrained from absolute quantitation of KLKs in CCM due to (a) the proof-of-concept nature of this data and (b) the strong impact of aspects such as cell lysis in SFM on the total protein content of CCM. For normalization purposes, we spiked in 1 μg avidin upon CCM harvesting to correct for potentially occurring technical variance during the sample preparation process. This data serves to illustrate which KLKs are amenable to robust detection by SRM in prototypical CCM of cultured pancreatic cancer cell lines. Of note, this data does not address limits of identification (LOI) or limits of quantitation (LOQ).

KLK6 was strongly detected with three peptides in the CCM of MiaPaCa-2 and the primary cell line PaCaDD-159. In other cell lines, KLK6 transitions remained below detection or were only marginally present. The inter-peptide differences of normalized peptide abundance, although originating from the same protein, probably derive from varying peptide-specific features like ionization or fragmentation (Bantscheff, Schirle, Sweetman, Rick, & Kuster, 2007).

KLK10 showed a different behavior and was detected with two peptides in the CCM of AsPC-1 and all primary cell lines, but not in MiaPaCa-2 (Fig. 1 - b). This is consistent with a previous report, which describes an elevated level of KLK10 in transcriptomic and proteomic data of AsPC-1 compared to MiaPaCa-2 (X. Y. Cao et al., 2018).

KLK3/PSA, which is clinically used as a biomarker for prostate cancer (Hong, 2014), was detected in all investigated pancreatic tumor cell lines (Fig. 1 - c). KLK5 and KLK7 were strongly detected in CCM of MiaPaCa-2 (Fig. 1 - d, e), which, together with KLK6, is consistent with a previous report that investigated KLK protein levels in CCM by ELISA (Shaw & Diamandis, 2008).

### KLK Detection in clinical FFPE tissue samples

Based on the robust, mass spectrometry-based detection of KLKs in CCM of cultured pancreatic cancer cells, we sought to investigate the presence of KLKs in pancreatic FFPE tissue samples using a targeted proteomics approach. We have refrained from absolute quantitation of KLKs in these samples since this data primarily serves to illustrate in which pancreatic pathological condition (PDAC, AMPAC, CP) KLK6, KLK7 or KLK10 are robustly detectable. This approach was motivated by KLKs having escaped detection by explorative mass spectrometry in previous pilot studies (data not shown). Of note, this data does not address LOI or LOQ.

We assembled a cohort including cases of PDAC (n=14), non-malignant adjacent pancreas (n=11), chronic pancreatitis (n=7), normal pancreas (n=5), and ampullary cancer (n=3). Basic patient information is summarized in Table S3. Tissue samples underwent targeted mass spectrometric analysis using parallel reaction monitoring (PRM) focusing on KLK6, KLK7, and KLK10, as these KLKs are considered promising diagnostic and/or prognostic candidates for pancreatic cancer (Candido et al., 2021) and showed robust detection in our previous cell culture experiment (Fig. 1).

Qualitative evaluation of PDAC tissue data revealed the near-consistent detection of KLK6 (14/17 cases) and KLK10 (15/17 cases) in the malignant tissue entities PDAC and AMPAC. In the entire set of non-malignant samples (total n=23) KLK6 and 10 were absent in all samples except for one case each (Fig. 2 - a). The obviously stronger presence of KLK6 and KLK10 in PDAC/AMPAC compared to CP or benign/normal pancreas is in accordance with previous publications (Candido et al., 2021; Hustinx et al., 2004; Ruckert et al., 2008; Srinivasan et al., 2022; Van Heek et al., 2004; Yousef, Borgono, Popalis, et al., 2004; Yousef, Borgono, White, et al., 2004).

**Fig. 2:**
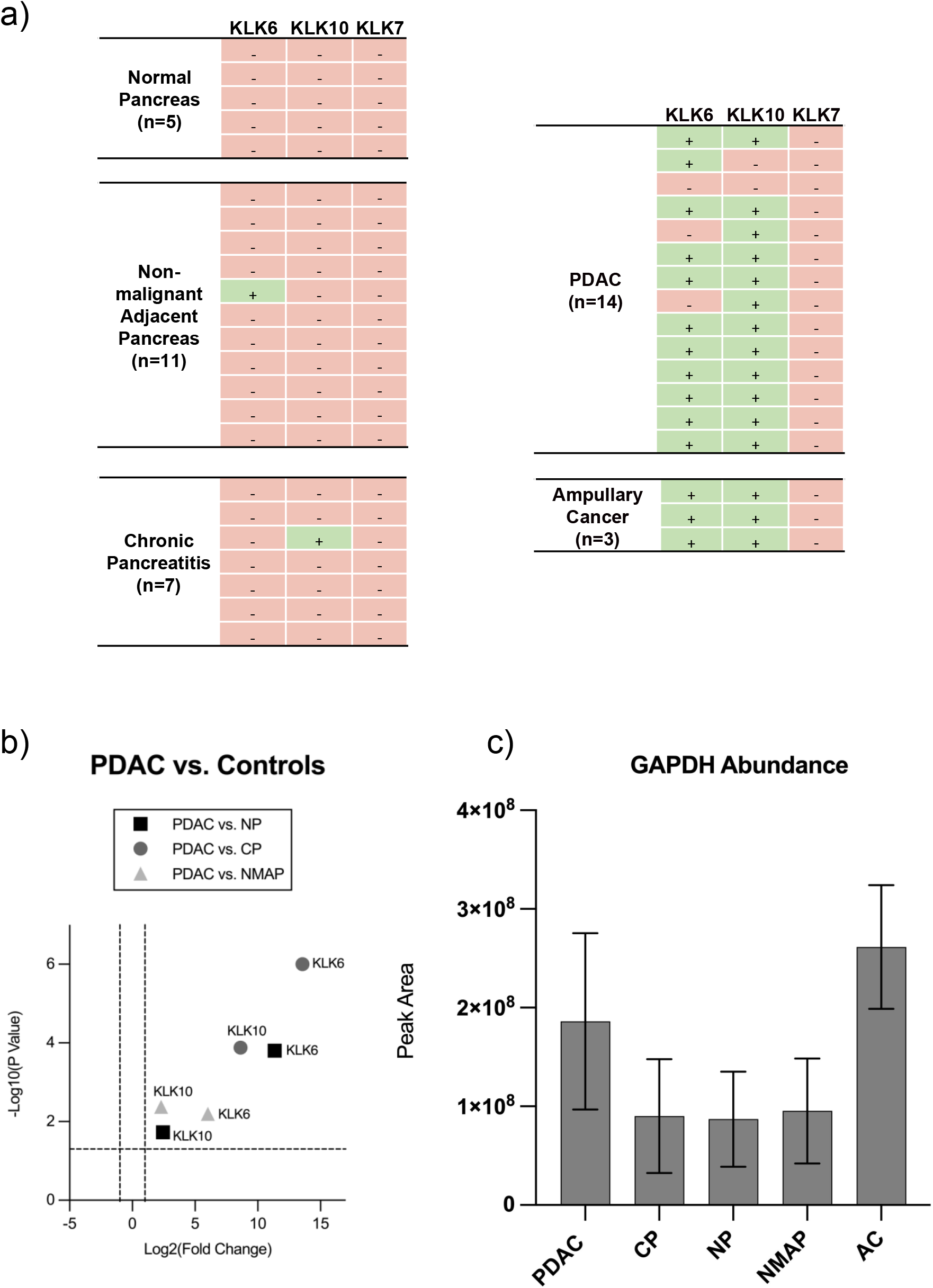
Qualitative and quantitative analysis of kallikrein proteins and GAPDH in clinical FFPE tissue samples by targeted mass spectrometry. Clinical FFPE tissue samples deriving from pancreatic ductal adenocarcinoma (PDAC), non-malignant adjacent pancreas (NMAP), chronic pancreatitis (CP), normal pancreas (NP) and ampullary cancer (AMPAC) were subjected to targeted mass spectrometric analysis using parallel reaction monitoring (PRM). Samples were measured in randomized sample order. (a) Focusing on KLK6, KLK7 and KLK10 enabled the qualitative evaluation of their detectable presence or absence in the respective tissue entities. (b) Results of pairwise quantitative comparisons (two-sample t-test) between PDAC and the respective controls (NP, CP and NMAP) are illustrated as a combined volcano plot depicting KLK6 and KLK10. The log2 fold changes are plotted on the x-axis and corresponding adjusted p-values in -log10 scale are shown on the y-axis. The applied log2 fold change cut-off was set to 2-fold, while the adjusted p-value cut-off was set to 0.05 (1.3 in -log10 scale), each depicted as dashed lines. A log2 fold change > 0 corresponds to an upregulation in the PDAC condition. (c) Evaluation of GAPDH-abundance (represented as peak area) in the respective tissue entities is shown as bar plot. Error bars correspond to standard deviation.

Although detected in our preliminary cell culture experiment (Fig. 1), KLK7 failed to be detected in any of the clinical FFPE tissue samples. This is in contrast to several publications that could show the presence of KLK7 in PDAC tissue either on transcriptomic level or immunodetection (Candido et al., 2021; Du et al., 2018; S. K. Johnson et al., 2007).

Although KLK6 and 10 remained below the LOD in benign pancreas (normal pancreas, non-malignant adjacent pancreas, and chronic pancreatitis), chromatographic background signals can be used to assess the extent of their upregulation in PDAC or AMPAC.

This semi-quantitative evaluation of KLK6 and KLK10 in PDAC compared to benign tissue controls showed a significant (p < 0.05, two-sample t-test using Benjamini Hochberg) upregulation of both kallikreins in PDAC compared to all controls (Fig. 2 - b). This is consistent with several previous reports (Candido et al., 2021; Yousef, Borgono, Popalis, et al., 2004) and has even been associated with shorter patient survival (Candido et al., 2021; Ruckert et al., 2008). In addition, KLK6 has been reported to be one of the most differentially expressed gene in pancreatic cancer compared with normal controls (Yousef, Borgono, Popalis, et al., 2004) which further supports our proteomic data, in which KLK6 revealed a more pronounced upregulation compared to KLK10. Furthermore, KLK6 and KLK10 also appeared to be significantly overexpressed in tissue samples of ampullary cancer compared to controls (Fig. S1), which underlines the already suggested similarities between pancreatic and ampullary cancer (Van Heek et al., 2004).

It should be noted that our bulk approach is not suited to distinguish intracellular (e.g. in secretory granules) and secreted KLKs, nor do we distinguish zymogens from activated KLKs. For the targeted proteomic array, we have included glyceraldehyde-3-phosphate dehydrogenase (GAPDH) as an endogenous protein for glycolytic energy metabolism. We observed varying abundance of GAPDH, with higher levels in malignant tissues (PDAC and AMPAC) and lower levels in benign tissues (Fig. 2 – c; raw peak areas from 1 μg injected tryptic sample). The variability of GAPDH abundance is in line with a study that found GAPDH to be among the genes with the highest expression variability in tumorous compared to normal pancreatic tissue (Rubie et al., 2005), possibly related to the “Warburg effect” in pancreatic cancer (Bose, Zhang, & Le, 2021; Warburg, 1924; Yang et al., 2020).

### Proteome Profiling allow Differentiation of different pancreatic entities

To obtain a global insight into the proteomic differences between the considered entities, an explorative analysis of clinical FFPE samples was additionally performed using LC-MS/MS operated in data independent acquisition (DIA) mode.

In a recent benchmarking study, we have shown that DIA-type proteomics is a powerful approach for characterizing the proteome biology of diseased or healthy tissue even in the presence of inter-individual heterogeneity (Fröhlich et al., 2022). Based on the benchmarking study, we opted for a gas-phase fractionation approach (Brian C. Searle et al., 2020) to generate an experiment-specific spectral library as well as using linear model of microarray analysis (limma) to statistically evaluate alterations in the proteome biology of the different entities.

Using the aforementioned approach, we identified a total of 5936 proteins in our cohort. On a per-sample basis, we identified and quantified 8073-31177 peptides and 2527-5023 proteins per sample (Fig. 3 – a). After median centering, the protein intensity distribution of the samples is comparable and spans several orders of magnitude (Fig. 3 - b). To further improve consistency within the dataset, we focused only on proteins with less than 20 % missingness per condition resulting in 2279 remaining proteins with a total of 2.1 % missing values in the overall dataset. The proteome coverage remains below the recently published CPTAC study on PDAC proteomics (L. Cao et al., 2021) but is substantially above the proteome coverage reported by non-fractionated DDA-type proteomics of PDAC (Oria et al., 2018).

**Fig. 3:**
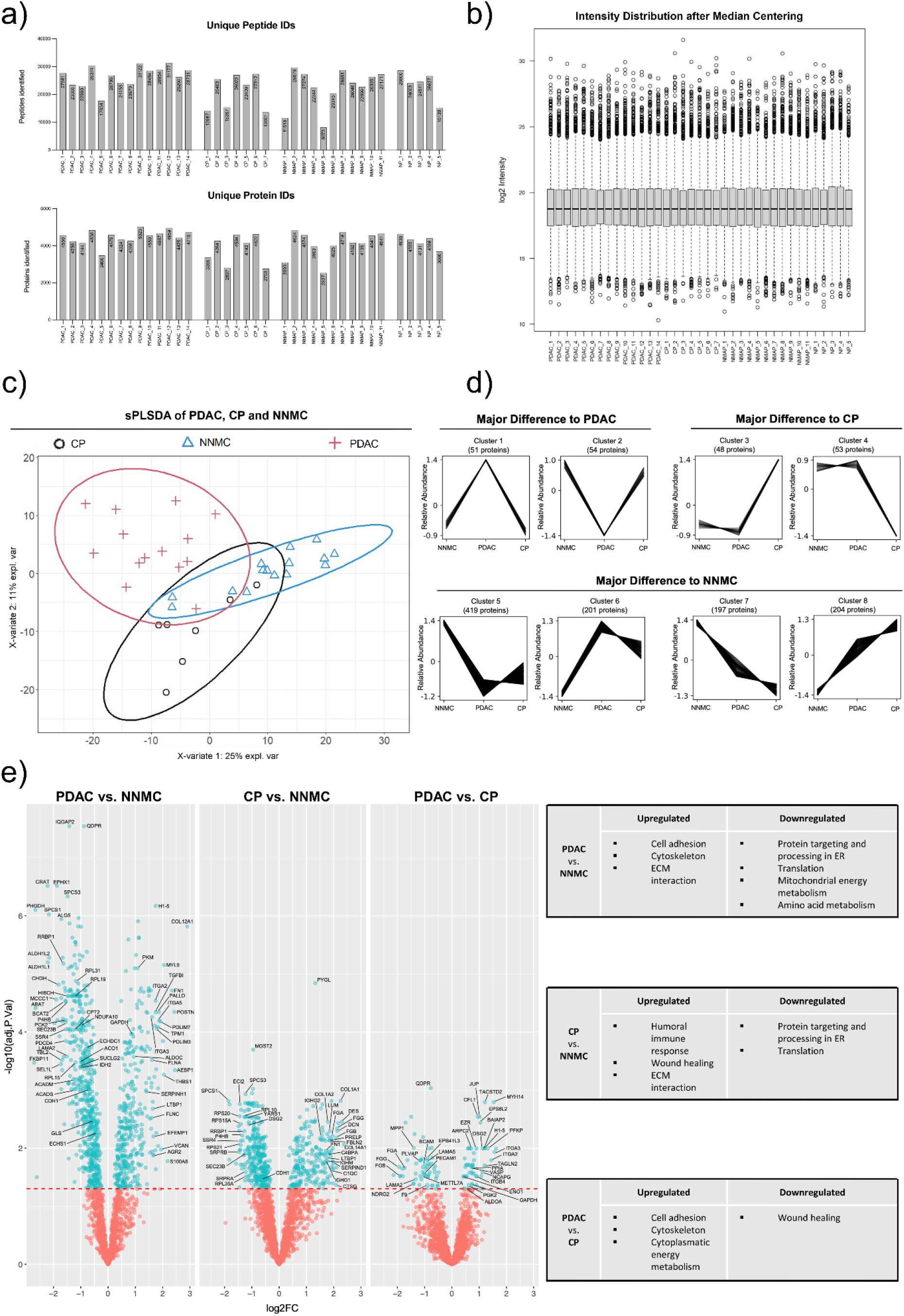
Proteomic profiling of different pancreatic tissue entities by explorative mass spectrometry. Clinical FFPE tissue samples deriving from pancreatic ductal adenocarcinoma (PDAC), non-malignant adjacent pancreas (NMAP), chronic pancreatitis (CP) and normal pancreas (NP) were subjected to explorative mass spectrometry analysis using data-independent acquisition (DIA) mode. Samples were measured in randomized sample order. (a) Resulting numbers of identified peptides and proteins are shown for each sample and tissue entity. (b) Box-and-Whisker-Plot shows the intensity distribution of each sample after median centering. In the following, the conditions NP and NMAP were combined to the condition normal non-malignant control (NNMC). (c) Protein profiles of each sample together with the corresponding condition annotation (CP, NNMC, PDAC) were submitted to sPLS-DA analysis. X- and y-axis represent the percentage of explained variance of the respective component, while ellipses represent the 95 % confidence intervals. (d) Depending on the measured abundance in different conditions (NNMC, PDAC, CP), the identified proteins were assigned to different co-abundance clusters with a confidence interval of 95 % by using the Clust algorithm. Each line represents an individual protein, while the y-axis illustrates the relative abundance change after Z-score normalization. Co-abundance clusters were sorted into three subgroups, depending on the conditions showing an abundance change compared to the normal control. The number of assigned proteins per cluster are shown above each graph. (e) For differential expression analysis, a pairwise multigroup limma approach was used to compare the conditions “PDAC vs. NNMC”, “CP vs. NNMC” and “PDAC vs. CP” while results were illustrated as volcano plots. The log2 fold changes (log2FC) are plotted on the x-axis and corresponding adjusted p-values in -log10 scale are shown on the y-axis. The applied adjusted p-value cutoff was set to 0.05 (1.3 in -log10 scale, depicted as horizontal line). Each plot highlights significantly up- or down-regulated proteins (blue). Hereby, a log2FC > 0 corresponds to an upregulation in the first-mentioned condition. Selected proteins are marked with their respective gene names. Gene Ontology (GO) enrichment analysis for the respective up- and down-regulated proteins were performed separately and a selection of the resulting significant GO terms (adjusted p-value < 0.05) are listed for each pairwise comparison.

We employed sparse partial least squares discriminant analysis (sPLS-DA) to depict the global proteomic divergence or similarity between the considered entities (Fig. S2). In this initial approach we observed near-complete overlap of the normal pancreas samples and the non-malignant adjacent pancreas samples (NMAP). Since highly similar proteomic profiles of normal and non-malignant adjacent pancreatic tissues have been reported previously (Cui et al., 2009), we merged these two conditions to a joint condition named normal non-malignant control (NNMC) for further analyses (Fig. S2). Further, we noticed that one NNMC sample was located beyond the 95 % confidence interval of NNMC (Fig. S3). Since this sample represented an NMAP sample with proximity to macrodissected tumor tissue, we consider the possibility of insufficient microdissection and actual presence of tumor tissue, leading to this control sample being considered inadequate and excluded from further analysis. Furthermore, the same sPLS-DA plot showed a complete overlap of the malign entities PDAC and ampullary cancer. Due to the low AMPAC case number (n=3), we refrained from further data analysis of the AMPAC proteomes.

The remaining dataset was submitted to a second sPLS-DA analysis to globally compare the proteome profiles of PDAC, CP, and NNMC (Fig. 3 – c). It is evident that all three entities show distinct clusters, suggesting proteomic differences, however with partial overlapping regions. Since the benign entities NNMC and CP have a slightly greater overlap than the malignant entity PDAC, one might speculate that the proteome difference between malignant and non-malignant entities appears to be more pronounced. The fact that benign pancreatic entities are crucially distinct from malignant tissues of PDAC is in agreement with the literature (Cui et al., 2009).

To identify protein groups exhibiting a comparable abundance behavior across the considered entities, we performed a co-abundance cluster analysis using the publicly available Clust algorithm (Abu-Jamous & Kelly, 2018). The clustering revealed eight distinct abundance-courses, with a total of 1227 assigned proteins (Fig. 3 - d). The eight clusters were further classified into three subgroups, depending on the condition showing the major abundance change. Considering the number of proteins assigned to each of these subgroups revealed the majority of proteins showing a similar abundance behavior of PDAC and CP compared to the control condition. This illustrates that both diseases introduce proteomic changes in the pancreas, emphasizing their deviation from the normal physiological state but also suggesting some joint characteristics (Vujasinovic et al., 2020). To find out whether certain biological processes or functions are particularly represented in the differently identified proteins, we further performed a gene ontology (GO) enrichment analysis (Fig. S5). While we observed a strong presence of proteins involved in cell adhesion and extracellular matrix (ECM) in PDAC, proteins involved in wound healing were evident in CP. In both PDAC and CP, there appears to be a neglect of normal exocrine pancreatic function.

In order to identify individual proteins that are significantly enriched or depleted in CP or PDAC compared to NNMC, we performed a multigroup limma approach for pairwise statistical testing to identify differentially expressed proteins in each entity. This analysis revealed 980, 588, and 112 significant hits (adjusted p-value ≤ 0.05) for the comparison PDAC vs. NNMC, CP vs. NNMC, and PDAC vs. CP, respectively (Fig. 3 – e, Sup.File1). To again evaluate whether certain biological processes or functions are especially represented within the differentially expressed proteins, we performed GO enrichment analysis for the up- and down-regulated proteins separately.

For the comparison of PDAC vs. NNMC, GO terms such as cytoskeleton, ECM interaction, and cell adhesion were found to be significantly upregulated in PDAC (Fig. 3 -e). Upregulated cell adhesion in PDAC compared to NNMC is also apparent when considering a heatmap representation of respectively associated proteins (Fig. S4 – a), which include e.g. fibronectin, vitronectin, thrombospondin, vinculin, fibulin, filamin, and several integrin subunits. Furthermore, our proteomic data also suggest an increase of cell adhesion-associated proteins in PDAC compared to CP, which could be also visualized via heatmap. For most epithelial tumors, downregulation of cell adhesion proteins is considered a hallmark feature. However, for PDAC, further studies support upregulation of cell adhesion proteins as compared to non-malignant control, e.g. for the tight junction protein ZO-1 (Kleeff et al., 2001) or the cell surface adhesion protein CEACAM6 (Pandey et al., 2019) as well as unique aspects of laminin and integrin biology (Humphries et al., 2022). To the best of our knowledge, the counter-intuitive enrichment of cell adhesion proteins in PDAC (as opposed to NNMC) has been largely under-appreciated.

A strong desmoplastic reaction and unique ECM biology is another hallmark feature of PDAC (Perez, Kearney, & Yeh, 2021). Our data recapitulates this aspect as PDAC revealed a significant upregulation of proteins associated with ECM interaction, which could be visualized via heatmap representation (Fig. S4 – b).

Downregulated GO terms in PDAC vs. NNMC comprise protein processing in the endoplasmic reticulum (ER) and mitochondrial energy metabolism (Fig. 3 - e). The former likely represents de-differentiation and loss of exocrine protein secretion functionality, whereas the latter may represent the Warburg effect. As such, glycolytic processes in the cytosol often appear to be upregulated, whereas oxidative phosphorylation in mitochondria is reduced, which has been also reported for pancreatic cancer (Yang et al., 2020). This effect is reflected by our data, as we observed a significant upregulation of several cytoplasmic glycolytic proteins (pyruvate kinase, PKM; fructose-bisphosphate aldolase C, ALDOC; ATP-dependent 6-phosphofructokinase, PFKP; phosphoglycerate kinase 2, PGK2; fructose-bisphosphate aldolase A, ALDOA; alpha-enolase, ENO1; and glyceraldehyde-3-phosphate dehydrogenase, GAPDH) in PDAC compared to the benign controls. Consistently, we also detected the decrease in the mitochondrial energy metabolism showing a significant downregulation of citrate cycle proteins (isocitrate dehydrogenase, IDH2; cytoplasmic aconitate hydratase, ACO1; succinate--CoA ligase, SUCLG2) in PDAC as well as an enriched GO term for downregulated mitochondrial energy metabolism. Furthermore, we detected a decreased amino acid degradation and downregulation of catabolic amino acid enzymes in PDAC, resulting in an increased availability of amino acids and thus providing an important source for cancer development and growth. Thereby, this data shows the metabolic reprogramming including the Warburg effect in PDAC on proteome level.

In chronic pancreatitis, GO terms like humoral immune response as well as wound healing were found to be significantly upregulated when compared to NNMC and show consistency with the literature (Bettac, Denk, Seufferlein, & Huber-Lang, 2017; Whittle & Hingorani, 2019). As representatives thereof, the complement protein c4b-binding protein alpha chain (C4BPA), several immunoglobulins (IGHG1, IGHG2, IGHM) and several fibrinogens (FGA, FGB, FGG) were found to be significantly upregulated in CP. Since the systemic activation of the complement system in pancreatitis is already known, our data further suggests the triggering of the complement system via the classical pathway as the C4BPA protein as well as immunoglobulins are known to be involved in this specific complement pathway (Bettac et al., 2017; Markiewski, Nilsson, Ekdahl, Mollnes, & Lambris, 2007). Thereby, our data confirms the inflammatory state of this entity on proteome level and highlights the role of the complement system in chronic pancreatitis.

Despite the described differences in the proteomes of PDAC and CP, we also identified several similarities when compared to NNMC. One of them is the common upregulation of ECM interaction and corresponding protein representatives including fibronectin (FN1), vinculin (VCL), fibulin (FBLN1), filamin (FLNA) and integrin subunits (e.g. ITGB1, ITGAV). The jointly increased abundance of proteins associated with ECM interaction in PDAC and CP is also apparent when considering a more global heatmap representation (Fig. S4 – b). This may reflect the role of ECM proteins in tissue remodelling not only during cancer progression but also during inflammatory processes (Shouji Shimoyama & Susanne Gansauge, 1995; Tian et al., 2019).

Furthermore, we observed a strong reduction of the exocrine pancreatic function in both PDAC and CP, since GO terms like translation and post-translational processing were found to be significantly downregulated as well as several representative proteins thereof including ribosomal proteins (RPL15, RPL19, RPL31, RPS20, RPS21, RPS15A, RPL10, RPL35A), members of protein export mechanisms (SPCS1, SPCS3, SRPRA, SRPRB) as well as protein processing (RRBP1, P4HB). This highlights the overall decreased gene expression in inflammatory as well as in malign pancreatic tissue.

Of note, we did not detect any KLK proteins in our DIA dataset. This may stem from technical limitations of the Q-Exactive plus mass spectrometer with regard to DIA proteomics, e.g. its limited scan speed. However, the same mass spectrometer was used for the successful PRM analysis of KLK6 and KLK10 described above, which is in line with earlier reports highlighting a much increased sensitivity of this instrumentation for targeted proteomics in the context of complex samples (Bauer et al., 2015). Hence, approaches that combine targeted and explorative methods may be particularly powerful (Hart-Smith, Reis, Waterhouse, & Wilkins, 2017).

### Pro-tumorigenic and tumor suppressor proteins in PDAC and CP

Sustaining proliferative signaling and evading growth suppressors are common hallmarks of cancer (Hanahan, 2022), which is why we specifically evaluated our proteomic dataset with regard to pro-tumorigenic and tumor suppressor proteins in PDAC and CP. As previously outlined, we have reduced our dataset to proteins with a maximum missingness of 20 % in PDAC, CP, and NNMC. This strict criterium might obscure tumor-driving or -suppressing proteins which are specifically present in PDAC and CP. Hence, we screened our proteomic data without missingness reduction for common protein representatives of each respective group using a reference list of Bosman *et al.* and further literature (Bosman, World Health Organization., & International Agency for Research on Cancer., 2010; Le et al., 2020). This resulted in the identification of ten pro-tumorigenic, and five tumor suppressor proteins in our data (Table 3 - a, b).

**Table 3:**
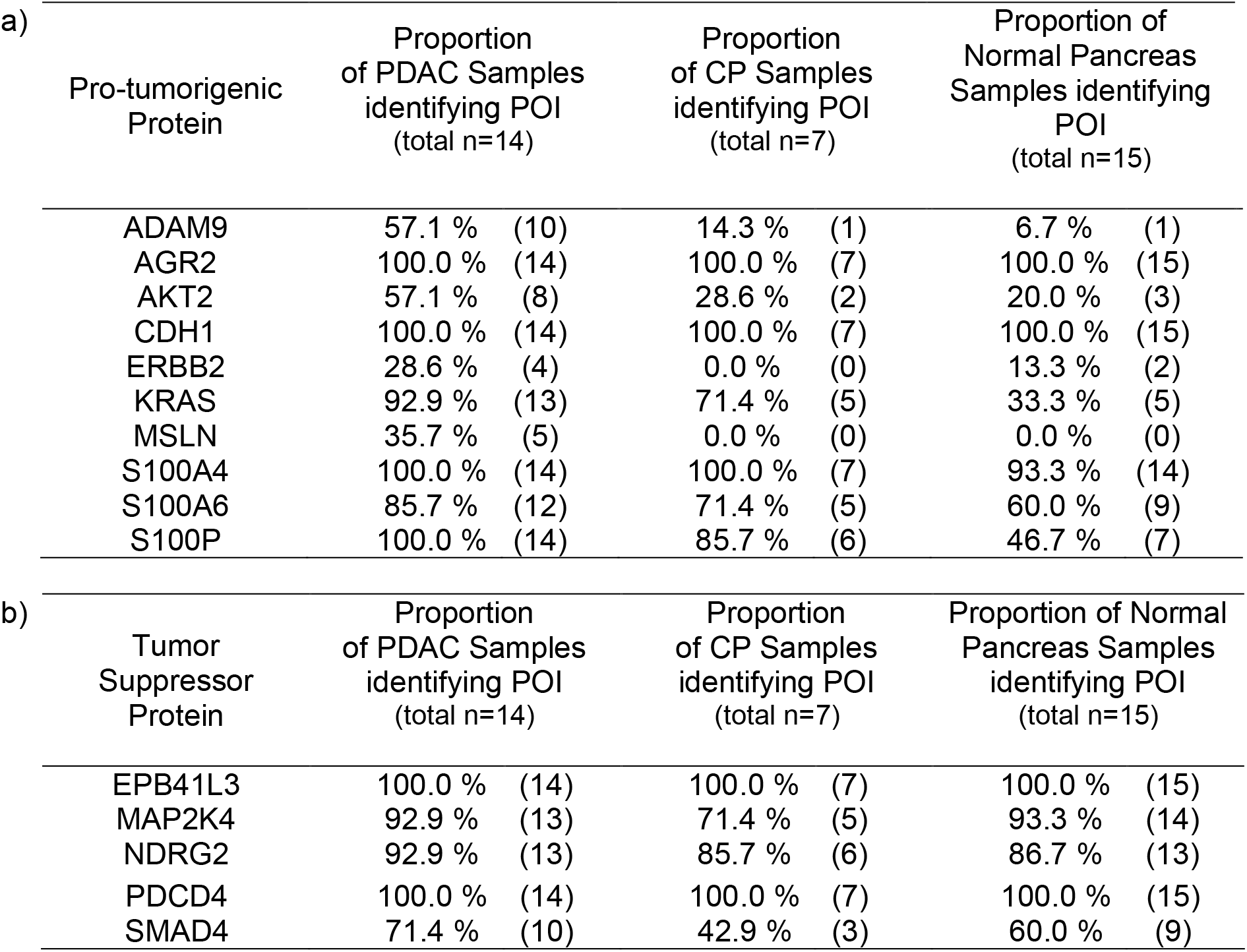
Number and proportion of identified pro-tumorigenic and tumor suppressor proteins in acquired explorative pancreatic dataset. The acquired dataset described and analyzed in Fig. 3 was screened for commonly known a) pro-tumorigenic and b) tumor suppressor proteins in tissues of PDAC, CP and Normal Pancreas. Listed values represent the percentage of samples in which the respective Protein Of Interest (POI) has been identified as well as the corresponding absolute number of samples in brackets.

We noticed enrichment in PDAC for the pro-tumorigenic proteins ADAM9, AKT2, ERBB2, KRAS, MSLN, S100A6 and S100P compared to normal pancreas samples (Table 3 – a). The strong presence of the cell surface protease ADAM9 in PDAC and its near-absence in normal pancreas samples is in line with earlier reports (Grutzmann et al., 2005). For ADAM9, we have previously shown its involvement in PDAC tumor biology by contributing to cell-matrix adhesion, anchorage independent growth, and neoangiogenesis (Oria et al., 2019). The proliferation-regulating protein KRAS has been described as the major driver gene in PDAC, which corroborates our proteomic findings (L. Cao et al., 2021).

Interestingly, CP shows an intermediate enrichment for ADAM9, AKT2, KRAS, S100A6 and S100P between PDAC and normal pancreas and thereby supports the hypothesis of CP being a preliminary stage to PDAC (Alhobayb et al., 2021; Del Poggetto et al., 2021; Vujasinovic et al., 2020). However, the receptor tyrosine kinase ERBB2 was not detected in CP and thereby showed an even diminished detection compared to normal pancreas samples.

Although it is expected to find the enrichment of pro-tumorigenic proteins in PDAC accompanied by the depletion of tumor suppressor proteins, we could not observe this for the considered tumor suppressor proteins in our proteomic data (Table 3 – b). Generally, tumor-driving proteins have lower missingness in PDAC and CP as opposed to NNMC while tumor-suppressing proteins have similar missingness in these entities (Table 3 - a, b).

### Semi-Tryptic Peptide Analysis To Estimate Endogenous Proteolytic Processing

We have previously shown that semi-tryptic (re-)analysis of proteomic data might yield novel insight into endogenous proteolytic processing prior to sample harvesting and eventual trypsination (Fahrner, Kook, Frohlich, Biniossek, & Schilling, 2021; Fretwurst et al., 2022; Shahinian et al., 2017). Naturally, this re-analysis can only shed light on non-tryptic, endogenous proteolytic processing. To further explore the acquired proteomic data, we performed a semi-tryptic peptide analysis to evaluate trypsin-independent endogenous proteolytic events and to compare them between the considered entities. Only semi-tryptic peptides were considered but not fully unspecific peptides. This revealed 3747 semi-tryptic peptides and corresponds to approximately 10 % of all peptides (35167 peptides) (data not shown). Comparing the intensity proportion of semi-tryptic peptides in relation to the overall intensity of all identified peptides revealed a significantly lower intensity-proportion in PDAC compared to the benign controls (Fig. 4 - a). Chronic pancreatitis showed the highest intensity proportion for semi-tryptic peptides, although its difference to normal pancreas was not significant. Likewise, when considering the absolute median log2 intensities of all semi-tryptic peptides per protein, we observed a slightly but significantly higher intensity for chronic pancreatitis although revealing less semi-tryptic peptides than PDAC and normal pancreas (Fig. 4 - b). This is expected and thereby validates previous literature as inflammatory tissue was previously described to reveal a higher proteolytic activity (Chakraborty & Bhattacharyya, 2013; Motta et al., 2021).

**Fig. 4:**
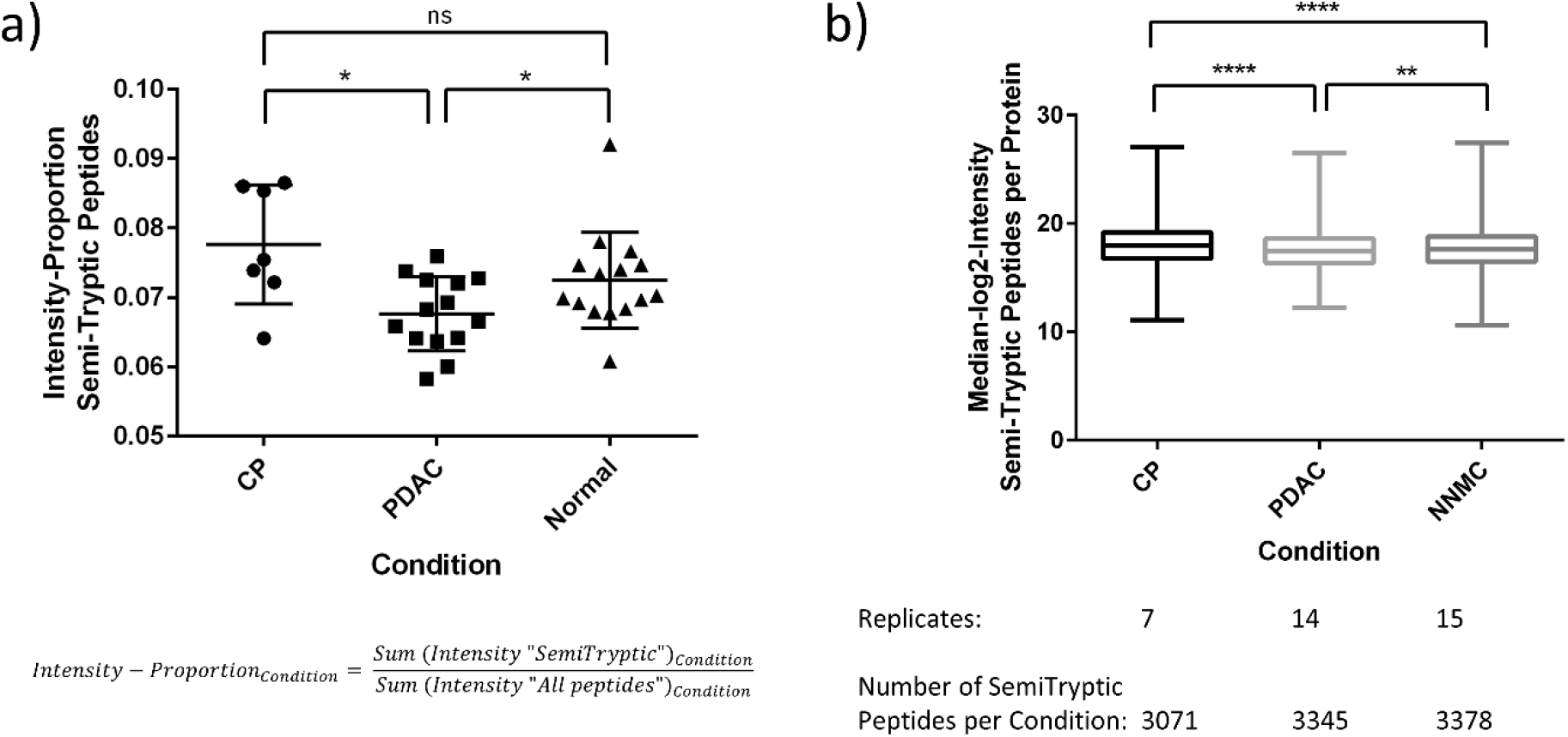
Semi-tryptic peptide analysis revealed different proteolytic activities in the considered entities. The acquired dataset described and analyzed in Fig. 3, was additionally analyzed with regard to semi-tryptic peptides. (a) Intensity proportion of semi-tryptic peptides in relation to all detected peptides is shown for each entity (CP, PDAC, Normal) as scatter plot with the horizontal line representing the mean and error bars corresponding to standard deviation. (b) Absolute median log2 intensities of all semi-tryptic peptides per protein were calculated for each entity (CP, PDAC, Normal) and are shown as Box-and-Whisker-Plot. For significance testing, a two-tailed Mann-Whitney U test was used because of unequal sample sizes between groups and not all groups passing normality tests (ns – non significant, *P ≤ 0.05, **P ≤ 0.01. ****P ≤ 0.0001).

### Proteogenomic Analysis and Identification of Single Amino Acid Variants

Proteogenomic analysis uses a combination of proteomics, genomics and/or transcriptomics data and enables the additional identification of potential sequence variants or copy number variations (CNVs) that could have not been detected when only considering a default human reference proteome (Nesvizhskii, 2014). This has recently been published for PDAC, where several correlations between CNVs and RNA or CNVs and protein level have been described (L. Cao et al., 2021). Although we do not have genomic data for our cohort and thereby cannot validate them as CNVs, we noticed that four differentially expressed proteins enriched in PDAC and two proteins depleted in PDAC were annotated as CNV in the aforementioned publication.

In order to identify PDAC- and CP-related single amino acid variants (SAAVs) in our data, proteogenomic analysis was performed on the acquired proteomic data together with publicly accessible transcriptomic data for PDAC (Table S4 – a for used SRA-identifier) (Mills et al., 2022) as a proof-of-concept study. This resulted in the median identification of 51, 67 and 53 SAAVs per CP, PDAC and NNMC sample, respectively (Fig. 5 – a). Although PDAC reveals a tendency for higher mutational burden/load, we do not see a significant difference by orders of magnitude in PDAC or CP in SAAV numbers, nor when comparing the intensity of the respective SAAV peptides (Fig. S6). This lack of differentiation may be due to the fact that we are not only considering mutational burden but also, to a large extent, naturally occurring allele variants which are also not part of the default proteome sequence database.

**Fig. 5:**
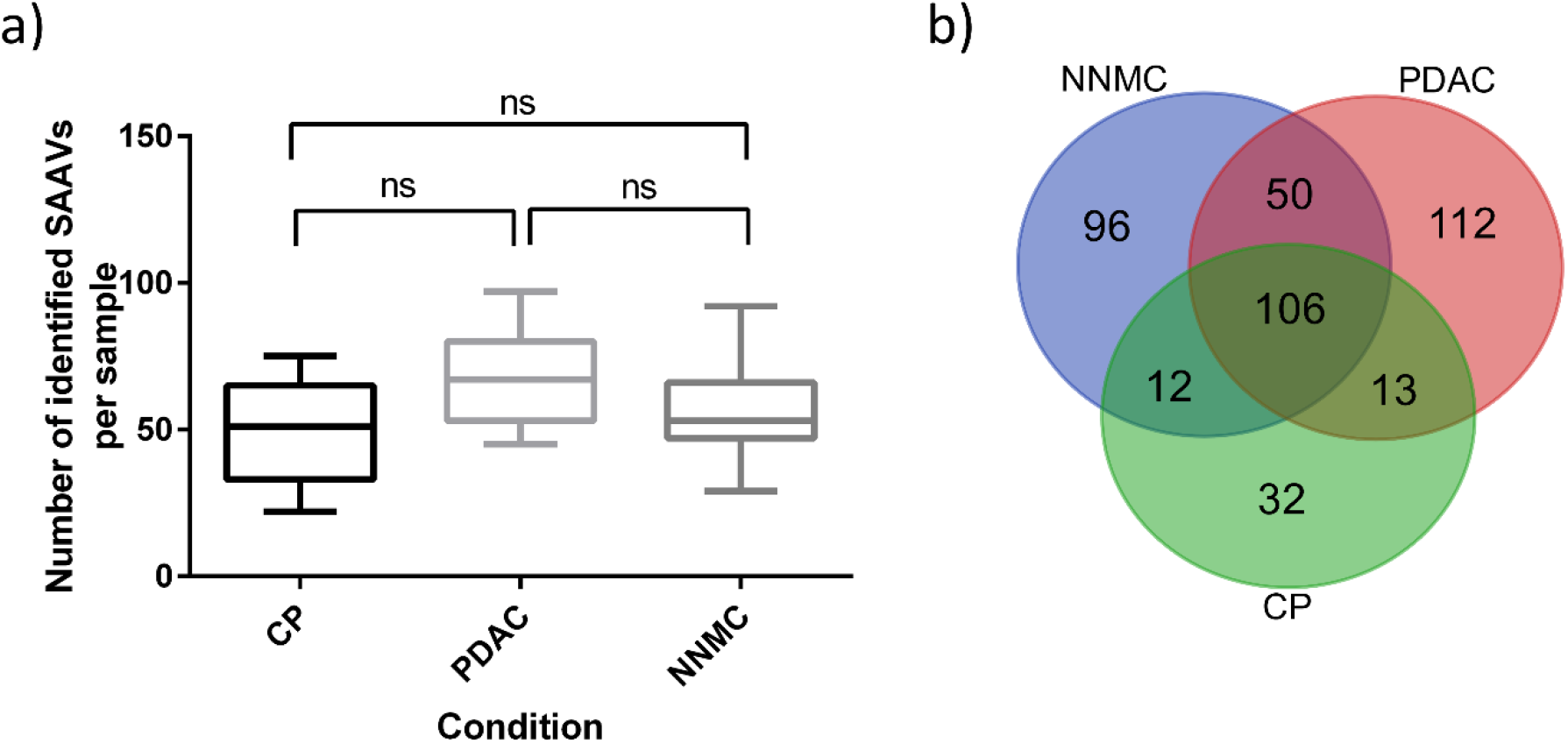
Proteogenomic analysis revealed single amino acid variants in PDAC and chronic pancreatitis. The acquired dataset described and analyzed in Fig. 3 was additionally analyzed with regard to single amino acid variants (SAAVs). (a) Number of identified SAAVs per sample is shown for each entity (CP, PDAC, Normal) as Box-and-Whisker plot. For statistical testing, one-way ANOVA with Tukey’s multiple comparisons test was used (ns – non significant). (b) The Venn diagram shows overlapping and specific SAAVs for each condition.

Comparing the identified SAAVs between the conditions, revealed 32 and 112 SAAVs, which specifically occur in CP and PDAC, respectively (Fig. 5 – b, Sup.File2). From the 112 PDAC-specific SAAVs, 16 were annotated as “pathogenic” (FATHMM-prediction) in the publicly available COSMIC database (Tate et al., 2018). Among these, the GTPase KRAS was found to be mutated in PDAC by the G12V substitution, which has been described to be one of the most frequently accumulated hotspots point mutations in PDAC (L. Cao et al., 2021; Topham et al., 2022) and thereby validates our findings. Of note, for CP, we found a S106L substitution of HRAS, which is another member of the RAS family comprising the most frequently mutated oncogene family in cancer. In addition, we found an E2410* nonsense mutation (prematurely terminated protein) of the Ankyrin repeat and KH domain-containing protein 1 (ANKHD1, UniprotID: Q8IWZ3) in PDAC, which native form has been suggested to be involved in cell proliferation of various cancers (Almeida & Machado-Neto, 2020; Fragiadaki & Zeidler, 2018). Furthermore, ANKHD1 has been described to inhibit Cyclin-dependent kinase inhibitor 1 (CDKN1A) expression in pancreatic cancer and is even suggested as potential therapeutic target (Ren, Sun, Li, Wu, & Jin, 2022). However, to evaluate the influence of the here observed SAAV, further investigations are required which is beyond the scope of this project.

## Conclusion

Our study suggests that KLK6 and 10 contribute to the tumor biology of pancreatic cancer due to their cell line specific secretion and significant upregulation in malignant entities such as PDAC and ampullary cancer compared to chronic pancreatitis and normal non-malignant pancreatic tissue. Explorative mass spectrometry clearly showed demarcated proteomes of malignant pancreatic tissue from protein expression patterns of the benign pancreas, among others illustrating the Warburg effect and a strong ECM fingerprint in pancreatic cancer. Furthermore, chronic pancreatitis was shown to reveal a higher endogenous proteolytic activity and proteogenomic analysis identified KRAS and ANKHD1 sequence variants. Our study provides novel insights into the proteomic characterization of PDAC and chronic pancreatitis and its association with kallikrein proteases, although larger patient cohorts will be required to validate our findings.

## Supporting information

MultigroupLimma-Results

Proteogenomic-Results

## Availability of data

All mass spectrometry proteomics datasets used and/or analyzed during this study are available online at the MassIVE repository (http://massive.ucsd.edu/; dataset identifier: MSV000090255)

## Ethics

This study was performed in line with the principles of the Declaration of Helsinki. Approval was granted by the Ethics Committee of Heidelberg University (02.01.2017 / 2012-293N-MA). Informed consent was obtained from all individual participants included in the study.

## Funding statement

OS acknowledges funding by the Deutsche Forschungsgemeinschaft (DFG, projects 446058856, 466359513, 444936968, 405351425, 431336276, 431984000 (SFB 1453 “Nephrogenetics”), 441891347 (SFB 1479 “OncoEscape”), 423813989 (GRK 2606 “ProtPath”), 322977937 (GRK 2344 “MeInBio”)), the ERA PerMed programme (BMBF, 01KU1916, 01KU1915A), the German-Israel Foundation (grant no. 1444), and the German Consortium for Translational Cancer Research (project Impro-Rec).

## Conflicts of Interest

The authors declare that they have no known competing financial interests or personal relationships that could have appeared to influence the work reported in this paper.

## Acknowledgement

The authors thank Bettina Wehrle for her technical support during the cell culture experiments.

## Author’s contributions

Janina Werner, Patrick Bernhard and Oliver Schilling contributed to the study conception and design. Janina Werner performed cell culture experiments and proteomic sample preparation. Janina Werner and Patrick Bernhard performed proteomic measurements. Patrick Bernhard analyzed the proteomic data. Janina Werner and Patrick Bernhard interpreted and visualized the proteomic data and drafted the manuscript. Matthias Fahrner established DIA-type measurements and supported during the data acquisition. Miguel Cosenza and Niko Pinter established various proteomic data analysis workflows and supported during their application. Prama Pallavi and Johannes Eberhard screened and provided patient samples and corresponding clinical data from Mannheim. Peter Bronsert screened and provided patient samples and corresponding clinical data from Biobank Freiburg. Felix Rückert and Oliver Schilling were major contributors in the revision of the manuscript. Oliver Schilling provided resources and funding. All authors read and approved the final manuscript.

## Supplementary Material

**Table S1:**
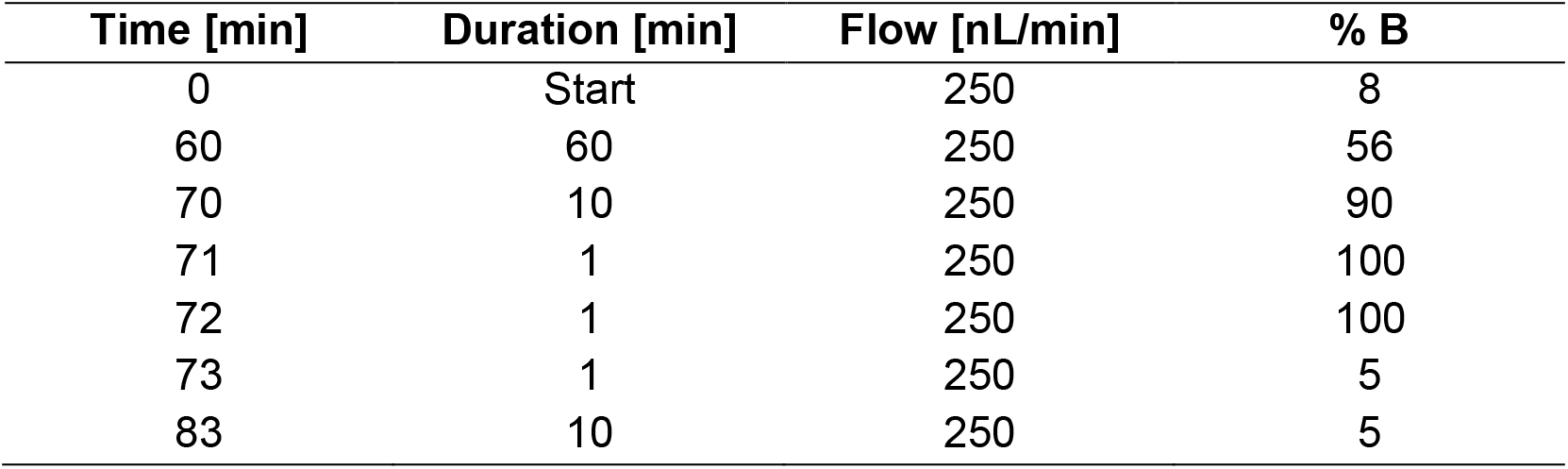
Applied chromatographic gradient for peptide separation prior to mass spectrometric measurement using SRM. The table describes the proportion of Buffer B (80 % v/v acetonitrile, 0.1 % v/v formic acid) in Buffer A (0.1 % v/v formic acid) over time together with the corresponding flow rate.

**Table S2:**
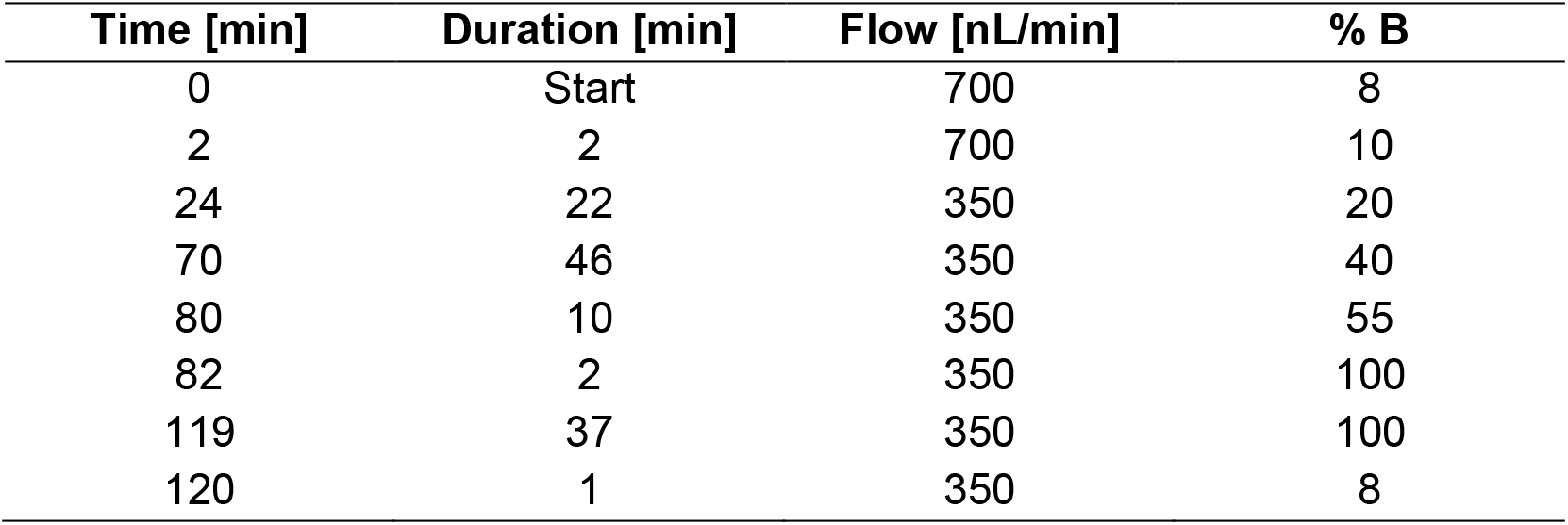
Applied chromatographic gradient for peptide separation prior to mass spectrometric measurement using PRM and DIA. The table describes the proportion of Buffer B (80 % v/v acetonitrile, 0.1 % v/v formic acid) in Buffer A (0.1 % v/v formic acid) over time together with the corresponding flow rate.

| Time [min] | Duration [min] | Flow [nL/min] | % B |
| --- | --- | --- | --- |
| 0 | Start | 700 | 8 |
| 2 | 2 | 700 | 10 |
| 24 | 22 | 350 | 20 |
| 70 | 46 | 350 | 40 |
| 80 | 10 | 350 | 55 |
| 82 | 2 | 350 | 100 |
| 119 | 37 | 350 | 100 |
| 120 | 1 | 350 | 8 |

**Table S3:** Cohort information of clinical FFPE tissue samples. The cohort contains tissues of PDAC (n=14), non-malignant adjacent pancreas (n=11), chronic pancreatitis (n=7), normal pancreas (n=5), and ampullary cancer (n=3). The table shows the sex of the patients, the age at the time of sampling, and the respective TNM classification.

| Entity | Sample Number | Sex | Age at Sampling | TNM-System & Grading |  |  |  |
| --- | --- | --- | --- | --- | --- | --- | --- |
|  |  |  |  | T | N | M | G |
| PDAC | 1 | M | 63 | 3 | 1 | 0 | 2 |
|  | 2 | F | 59 | 2 | 1 | 0 | 3 |
|  | 3 | M | 59 | 1 | 0 | 0 | 3 |
|  | 4 | M | 57 | 2 | 1 | 0 | 3 |
|  | 5 | M | 65 | 2 | 2 | 0 | 2 |
|  | 6 | M | 79 | 2 | 0 | 0 | 3 |
|  | 7 | M | 59 | 3 | 2 | 0 | 2 |
|  | 8 | F | 52 | 1 | 0 | 0 | 2 |
|  | 9 | F | 64 | 2 | 1 | 0 | - |
|  | 10 | M | 49 | 4 | 2 | 0 | 2 |
|  | 11 | M | 78 | 2 | 0 | 0 | 3 |
|  | 12 | F | 49 | - | 0 | - | 1 |
|  | 13 | - | - | - | - | - | - |
|  | 14 | F | 60 | 4 | - | 1 | - |
| CP | 1 | M | 57 | Not applicable |  |  |  |
|  | 2 | F | 48 | Not applicable |  |  |  |
|  | 3 | F | 63 | Not applicable |  |  |  |
|  | 4 | M | 52 | Not applicable |  |  |  |
|  | 5 | F | 77 | Not applicable |  |  |  |
|  | 6 | M | 40 | Not applicable |  |  |  |
|  | 7 | M | 71 | Not applicable |  |  |  |
| Ampullary Cancer | 1 | M | 82 | 2 | 0 | 0 | - |
|  | 2 | F | 62 | 3 | 1 | 0 | 2 |
|  | 3 | M | 62 | 3 | 2 | 0 | 3 |
| Normal Pancreas | 1 | - | 80 | Not applicable |  |  |  |
|  | 2 | - | 43 | Not applicable |  |  |  |
|  | 3 | - | 46 | Not applicable |  |  |  |
|  | 4 | - | 67 | Not applicable |  |  |  |
|  | 5 | - | 59 | Not applicable |  |  |  |

**Fig. S1:**
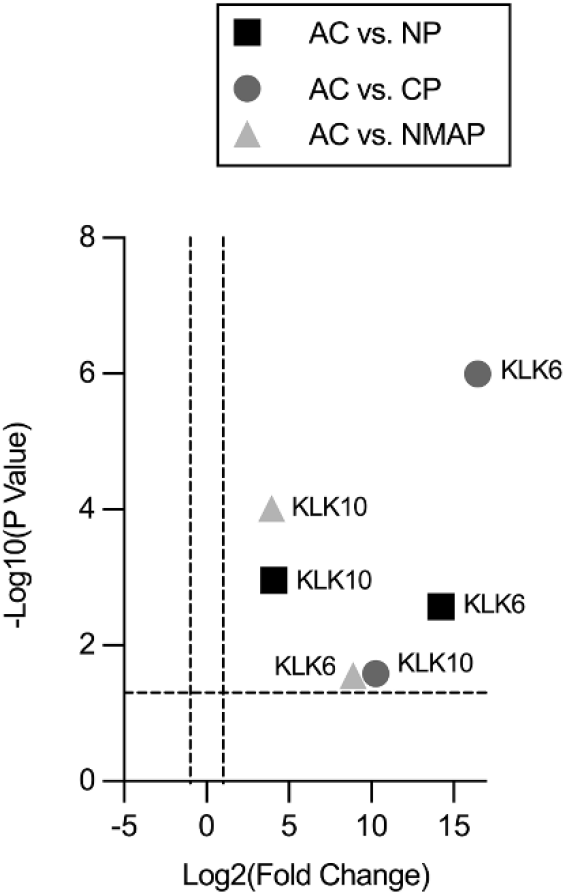
Pairwise Quantitative Comparisons of Kallikrein 6 and 10 Protein Levels between Ampullary Cancer and the respective Controls. Clinical FFPE tissue samples deriving from non-malignant adjacent pancreas (NMAP), chronic pancreatitis (CP), normal pancreas (NP) and ampullary cancer (AMPAC) were subjected to targeted mass spectrometric analysis using parallel reaction monitoring (PRM). Samples were measured in randomized sample order. Results are illustrated as combined volcano plot depicting KLK6 and KLK10. The log2 fold changes are plotted on the x-axis and corresponding adjusted p-values in -log10 scale are shown on the y-axis. The applied log2 fold change cutt-off was set to 2-fold, while the adjusted p-value cut-off was set to 0.05 (1.3 in -log10 scale), each depicted as dashed lines. A log2 fold change > 0 corresponds to an upregulation in the PDAC condition.

**Fig. S2:**
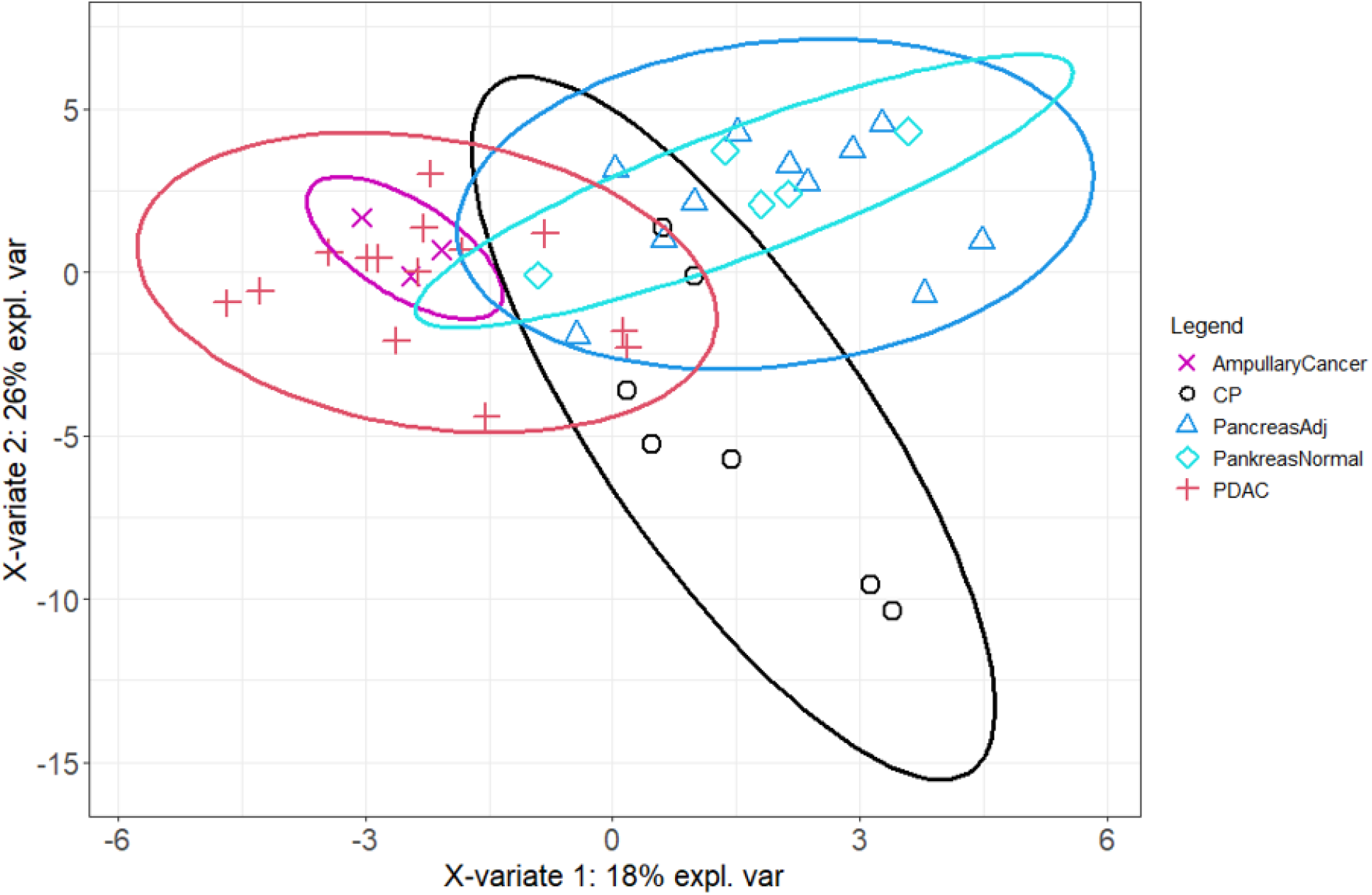
Sparse Least Squares Discriminant Analysis (sPLS-DA) of DIA data with original condition groups. Protein profiles of each sample together with the corresponding original condition annotation (ampullary Cancer, chronic pancreatitis - CP, non-malignant adjacent pancreas-NMAP, normal pancreas, PDAC) were submitted to sPLS-DA analysis. X- and y-axis represent the percentage of explained variance of the respective component, while ellipses represent the 95 % confidence intervals.

**Fig. S3:**
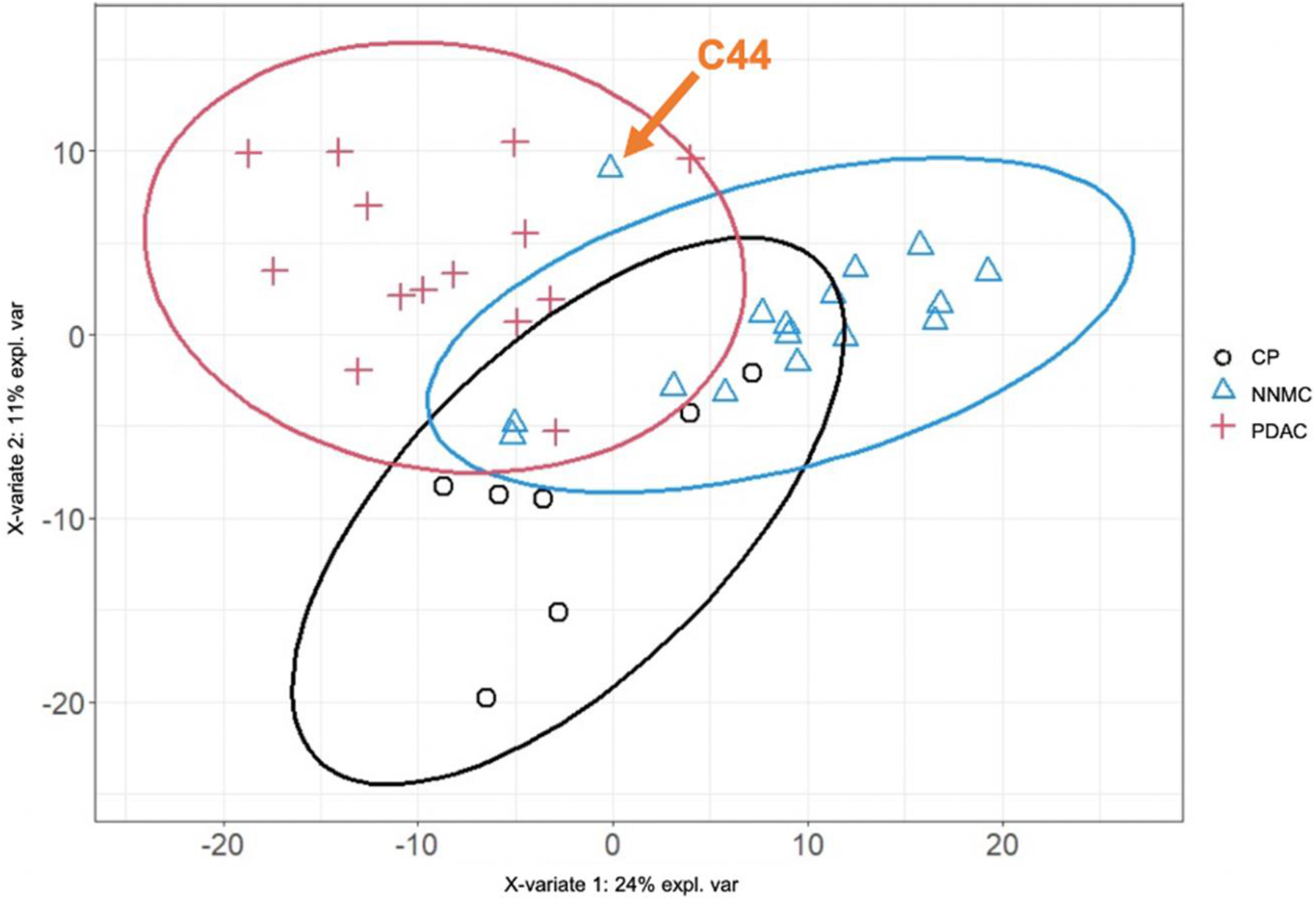
Sparse Least Squares Discriminant Analysis (sPLS-DA) of DIA data with adjusted condition groups. Protein profiles of each sample together with the corresponding original condition annotation (chronic pancreatitis - CP, normal non-malignant control - NNMC, PDAC) were submitted to sPLS-DA analysis. X- and y-axis represent the percentage of explained variance of the respective component, while ellipses represent the 95 % confidence intervals. The sample C44 is located outside the 95 % confidence interval for the NNMC condition (orange arrow).

**Fig. S4:**
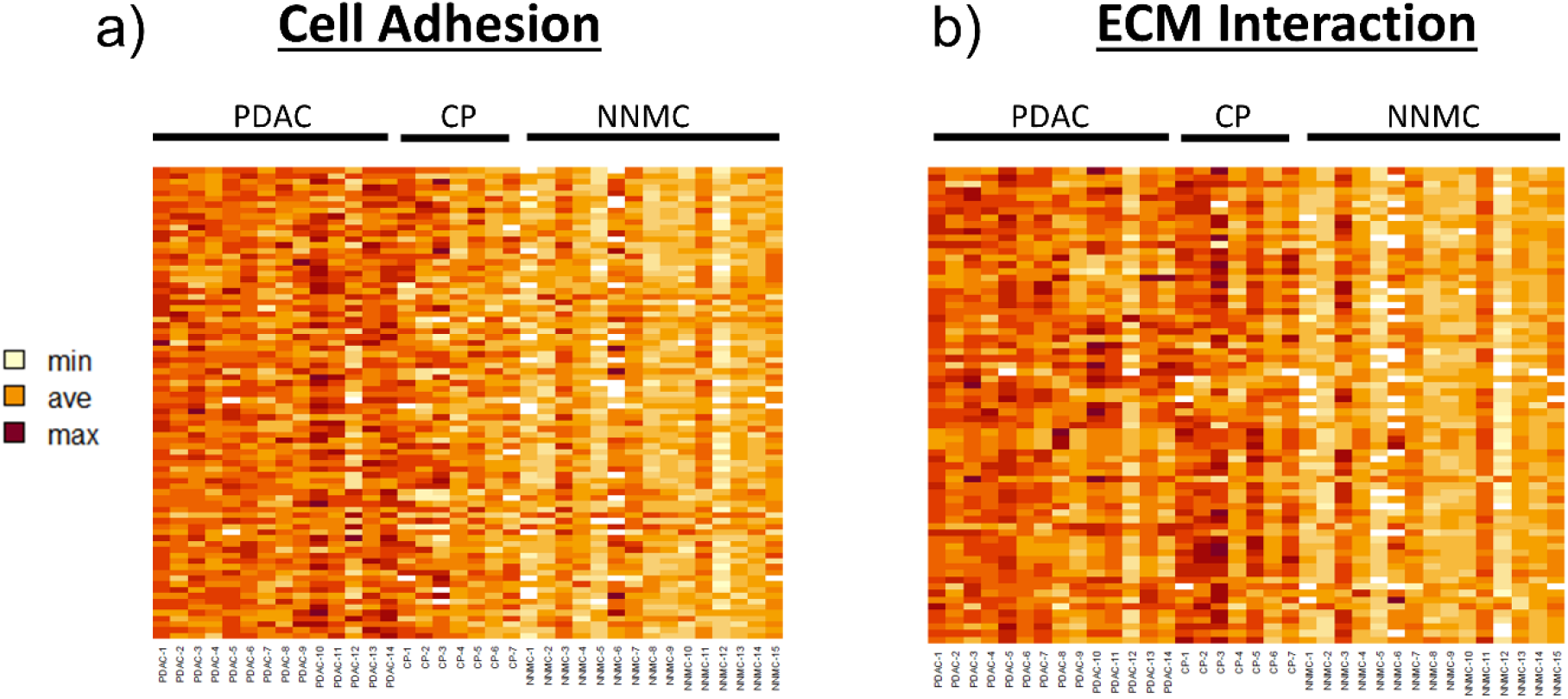
Heatmap Representation of Expression Profiles for Proteins associated with the GO-terms Cell Adhesion or ECM Interaction. Acquired proteomic dataset including samples from pancreatic ductal adenocarcinoma (PDAC), chronic pancreatitis (CP) or normal non-malignant control (NNMC) tissue was filtered for proteins associated with the gene ontology terms “Cell Adhesion” or “ECM Interaction”. Respective intensity values, which correspond to their abundance, was visualized via heatmap representing proteins as rows, conditions (PDAC, CP, NNMC) as columns and the respective protein abundance as colour-coded field. Samples from similar conditions were grouped together and indicated accordingly above the heatmap.

**Fig. S5:**
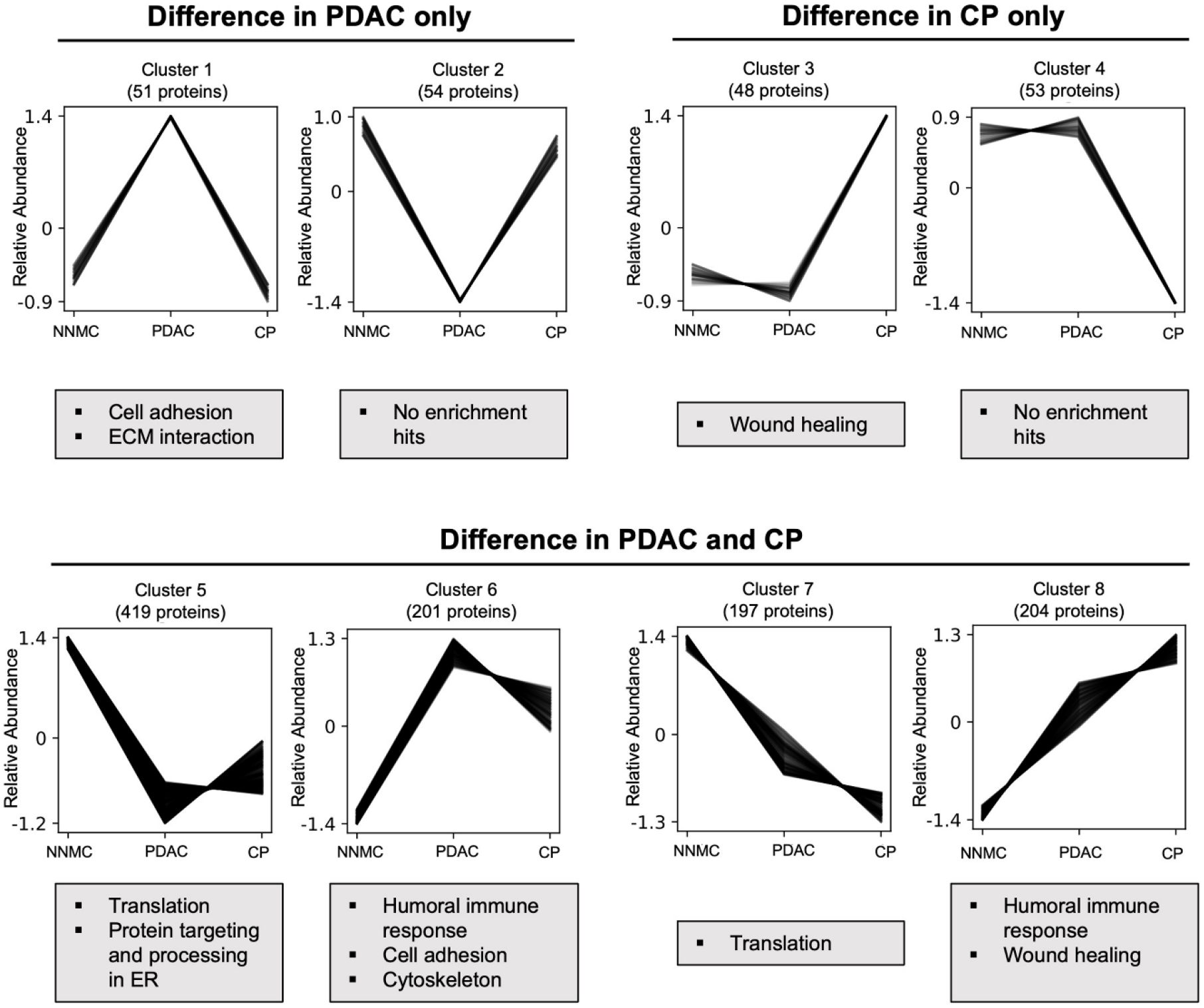
GO enrichment analysis of the co-abundance clusters. The eight clusters resulting from the co-abundance cluster analysis were further subjected to a GO enrichment analysis. The respective significant GO terms (adjusted p-value < 0.05) are shown in the boxes below each cluster diagram.

**Table S4:**
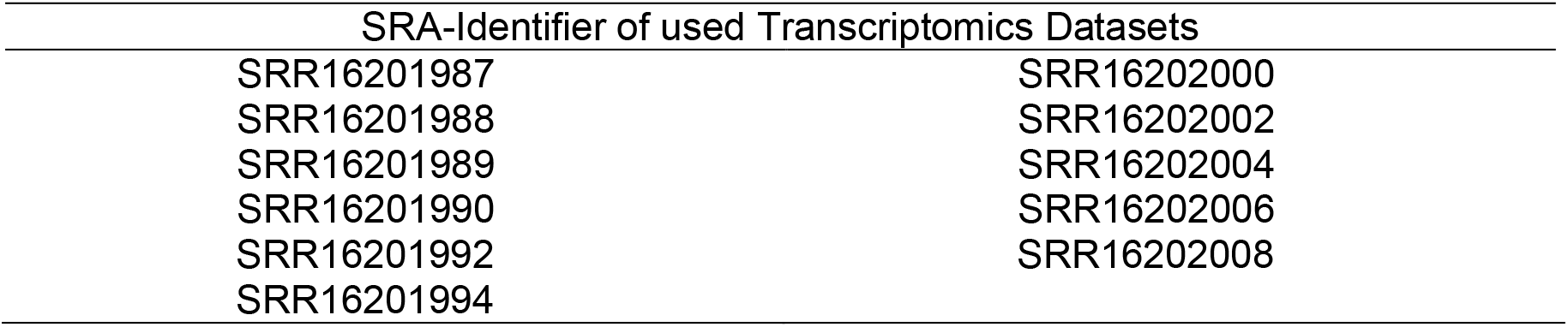
Used transcriptomics data for proteogenomic analysis. Publicly available transcriptomics data for PDAC (Mills et al., 2022) was used for proteogenomic analysis of acquired proteomics data. Corresponding Sequence Read Archive (SRA) identifier from NCBI are listed.

**Fig. S6:**
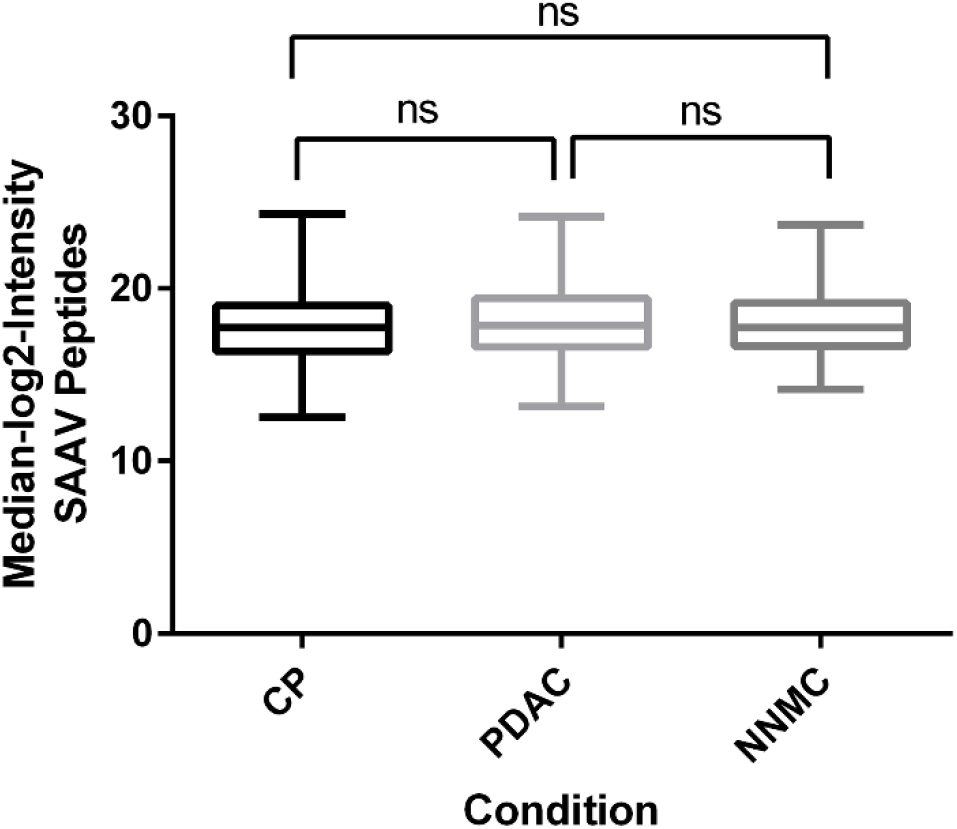
Peptide intensity comparison of identified single amino acid variants. Proteogenomic analysis revealed single amino acid variants (SAAVs) in the acquired dataset. Median log2 intensities of respectively identified SAAV peptides were calculated for each condition (CP, PDAC, NNMC) and are shown as Box-and-Whisker-Plot. For significance testing, one-way ANOVA with Tukey’s multiple comparisons test was used (ns – non significant).

## Abbreviations

AcLS: Acid-labile surfactant
ACN: Acetonitrile
Adj.P.Val: Adjusted p-value
AGC: Automatic Gain Control
BCA: Bicinchoninic Acid
CCM: Cell-Conditioned Medium
CP: Chronic Pancreatitis
DDA: Data Dependent Acquisition
DIA: Data Independent Acquisition
DMEM: Dulbecco’s Modified Eagle’s Medium
DTT: Dithiothreitol
EDTA: Ethylenediaminetetraacetic acid
EMT: Epithelial-Mesenchymal Transition
ESI: Electrospray Ionization
FA: Formic Acid
FC: Fold Change
FDR: False Discovery Rate
FFPE: Formalin-fixed paraffin-embedded
GO: Gene Ontology
HEPES: 4-(2-hydroxyethyl)-1-piperazineethanesulfonic acid
ID: Identification
iRT: Indexed Retention Time
KLK: kallikrein
LC: Liquid Chromatography
LysC: Lysyl Endopeptidase
MS: Mass Spectrometry
m/z: Mass-to-Charge Ratio
NMAP: non-malignant adjacent pancreas
NNMC: normal non-malignant control
PDAC: Pancreatic ductal adenocarcinoma
PRM: Parallel Reaction Monitoring
RPMI: Roswell Park Memorial Institute Medium
RT: Room Temperature
SAAV: Single Amino Acid Variation
Sp3: Single-Pot, Solid-Phase-Enhanced Sample-Preparation
sPLS-DA: sparse Partial Least Squares Discriminant Analysis
SRM: Selected Reaction Monitoring
TFA: Trifluoroacetic acid

## Notes

https://massive.ucsd.edu/ProteoSAFe/dataset.jsp?task=8683e4b630fa446f88d09730319115e6

### Competing Interest Statement

The authors have declared no competing interest.

